# Biophysical and pharmacokinetic characterization of a small-molecule inhibitor of RUNX1/ETO tetramerization with anti-leukemic effects

**DOI:** 10.1101/2021.12.22.473911

**Authors:** Mohanraj Gopalswamy, Tobias Kroeger, David Bickel, Benedikt Frieg, Shahina Akter, Stephan Schott-Verdugo, Aldino Viegas, Thomas Pauly, Manuela Mayer, Julia Przibilla, Jens Reiners, Luitgard Nagel-Steger, Sander H.J. Smits, Georg Groth, Manuel Etzkorn, Holger Gohlke

## Abstract

Acute myeloid leukemia (AML) is a malignant disease of immature myeloid cells and the most prevalent acute leukemia among adults. The oncogenic homo-tetrameric fusion protein RUNX1/ETO results from the chromosomal translocation t(8;21) and is found in AML patients. The nervy homology region 2 (NHR2) domain of ETO mediates tetramerization; this oligomerization is essential for oncogenic activity. Previously, we identified the first-in-class small-molecule inhibitor of NHR2 tetramer formation, **7.44**, which was shown to specifically interfere with NHR2, restore gene expression down-regulated by RUNX1/ETO, inhibit the proliferation of RUNX1/ETO-depending SKNO-1 cells, and reduce the RUNX1/ETO-related tumor growth in a mouse model. However, no biophysical and structural characterization of **7.44** binding to the NHR2 domain has been reported. Likewise, the compound has not been characterized as to physicochemical, pharmacokinetic, and toxicological properties. Here, we characterize the interaction between the NHR2 domain of RUNX1/ETO and **7.44** by biophysical assays and show that **7.44** interferes with NHR2 tetramer stability and leads to an increase in the dimer population of NHR2. The affinity of **7.44** with respect to binding to NHR2 is *K*_lig_ = 3.95 ± 1.28 μM. By NMR spectroscopy combined with molecular dynamics simulations, we show that **7.44** binds with both heteroaromatic moieties to NHR2 and interacts with or leads to conformational changes in the N-termini of the NHR2 tetramer. Finally, we demonstrate that **7.44** has favorable physicochemical, pharmacokinetic, and toxicological properties. Together with biochemical, cellular, and in vivo assessments, the results reveal **7.44** as a lead for further optimization towards targeted therapy of t(8;21) AML.

## Introduction

RUNX1/ETO is an oncogenic homotetrameric fusion protein found in t(8;21)-dependent acute myeloid leukemia (AML) patients [1]. This form of AML is characterized by an achieved complete remission for ~50% to 75% of patients [2], a relapse-free survival rate of ~20% to 30% [2], and a 5-year overall survival rate of ~50% [3]. Because no targeted treatment of t(8;21)-dependent AML is available at the moment, the discovery of new therapeutic targets with the potential to improve the overall survival rate is of utmost importance [4]. RUNX1/ETO is composed by the DNA-binding Runt-domain [5], the product of the RUNX1 gene, and by four nervy homology regions (NHR1-4), the product of the ETO gene (**Figure 1A**) [6]. The NHR2 domain is responsible for the tetramerization of RUNX1/ETO [7]. Furthermore, the NHR3- and NHR4-deleted versions of RUNX1/ETO and other short variants rapidly induce leukemia in a mouse bone marrow transplantation model [8]. Recently, the specificity of the NHR2 domain was displayed by replacing the NHR2 domain from full-length RUNX1/ETO with a structurally similar BCR tetramerization domain resulting in a loss of CD34+ progenitor cell expansion [9]. Thus, the NHR2-mediated tetramerization is necessary for the onset and maintenance of t(8;21)-dependent AML [10]. Hence, the disruption of RUNX1/ETO tetramerization is a promising strategy to fight AML.

**Figure 1.**
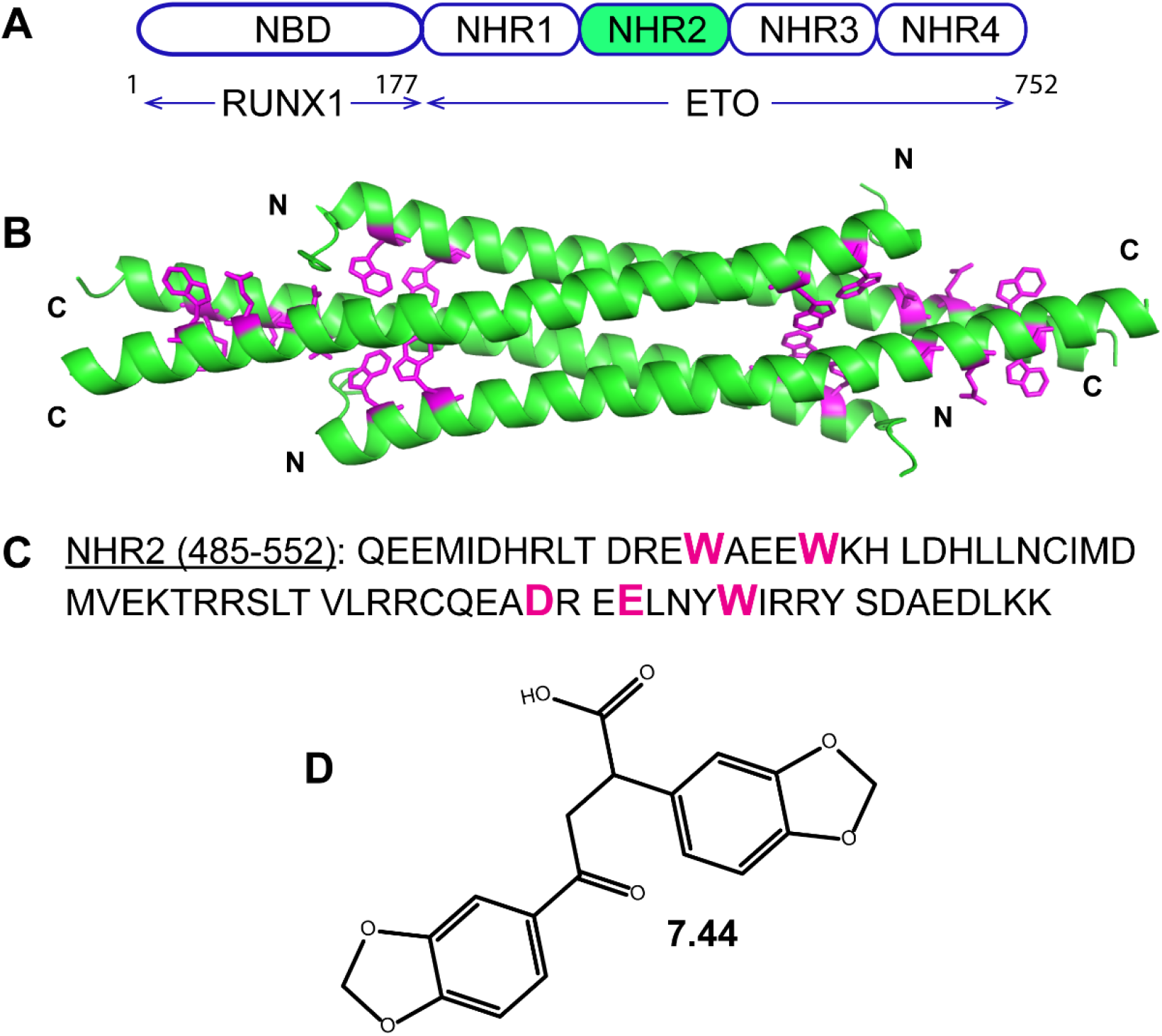
Schematic representation of RUNX1/ETO tetramerization. A) Depiction of the chimeric RUNX1/ETO fusion protein containing the nucleotide-binding domain (NBD) and four *nervy homology regions* (NHR). The NHR2 domain (marked in green) mediates homo-tetramerization. B) X-ray crystal structure (PDB ID 1WQ6) of the NHR2 domain. C) Amino acid sequence of the NHR2 domain. In B) and C) predicted hot spot residues [11] are highlighted in magenta. D) Molecular structure of **7.44**.

Previous studies identified five hot spots residues, W498, W502, D533, E536, and W540 **(Figure 1B and C**), as essential for the tetramerization NHR2 [11]. By mutating the hot spots to alanine, resulting in what has been termed M5 variant, we demonstrated that RUNX1/ETO-dimers, in contrast to tetramers, do not block myeloid differentiation and fail to induce AML in mice [10]. Thus, the hot spot residues of NHR2 represented a target for developing inhibitors of RUNX1/ETO tetramerization and a molecular intervention in t(8;21)-dependent AML that had not yet been addressed. An 18-mer peptide containing the hot spot residues inhibited the tetramerization of NHR2 with an *IC*_50_ of 250 μM (BS^3^ cross-linking assay) and 390 μM (ELISA) [11]. Although this example shows that inhibiting NHR2 tetramerization on a molecular level is possible, the 18-mer has unfavorable pharmacological properties. Therefore, exploiting the pattern of a subset of the hot spots (D533, E536, and W540) as a query for virtual screening, we identified the first-in-class small-molecule inhibitor of NHR2 tetramer formation, **7.44 (Figure 1D)** [11]. **7.44** was shown to specifically interfere with NHR2, restore gene expression down-regulated by RUNX1/ETO, inhibit the proliferation of RUNX1/ETO-depending SKNO-1 cells, and reduce the RUNX1/ETO-related tumor growth in a mouse model [12]. These findings, together with other favorable properties of **7.44** [11], such as low molecular weight, a high ligand efficiency, and a non-complex chemical structure, suggest that **7.44** could serve as a lead structure to guide the development of structurally related compounds with increased binding affinity, improved bioavailability, and enhanced anti-leukemic effects to inhibit RUNX1/ETO oncogenic function in t(8;21) AML. However, biophysical and structural characterization of **7.44** binding to the NHR2 domain has not been done yet. Likewise, the compound has not yet been characterized as to physicochemical, pharmacokinetic, and toxicological properties.

In this study, we established biophysical assays based on differential scanning fluorimetry (also termed thermal shift assay (TSA)) and microscale thermophoresis (MST) to determine the binding affinity of **7.44** to NHR2 and show that the dissociation constant of **7.44** from the NHR2 dimer, *K*_lig_, is 3.95 ± 1.28 μM. We also suggest how **7.44** interferes with NHR2 tetramer stability based on integrative modeling using saturation transfer difference NMR (STD-NMR) and multidimensional NMR experiments as well as molecular dynamics (MD) simulations of free ligand diffusion. These studies indicate that **7.44** binds at the NHR2 termini and reveal moieties of **7.44** important for binding. Finally, we report favorable physicochemical, pharmacokinetic, and toxicological properties of **7.44**. These results are a significant step towards understanding how **7.44** inhibits NHR2 tetramerization at the molecular level and should aid in the development of improved inhibitors of NHR2 tetramerization.

## Materials & Methods

### Cloning, expression, and purification

Codon-optimized synthetic DNA (Genscript Biotech) of the NHR2 domain (residues 485-552 of human RUNX1/ETO) or M5 variant (five residues of the NHR2 domain W498, W502, D533, E536, and W540 were mutated to alanine) was cloned into expression vector pETSUMO in *E. coli* Rosetta (DE3)pLysS (Invitrogen).

The NHR2 domain or M5 variant was expressed as a SUMO (small ubiquitin-related modifier) fusion protein. The transformed *E. coli* was transferred into dYT-medium (10 g yeast extract, 5 g NaCl, and 16 g tryptone or peptone), containing kanamycin and chloramphenicol, and incubated at 37 °C until an OD_600_ of ~1 was reached. The expression was induced by adding isopropyl-1-thio-β-D-galactopyranoside to 1 mM final concentration. The expression was continued overnight at 25 °C. The cells were harvested by centrifugation and disrupted via sonification.

The same purification protocol for the SUMO fusion protein was used as mentioned earlier [13]. In brief, the fusion protein was purified by nickel affinity chromatography (cell lysis buffer-50 mM sodium phosphate, 300 mM sodium chloride, 10 mM imidazole, pH 8) on a HisTrap column (elusion buffer-50 mM sodium phosphate, 300 mM sodium chloride, 500 mM imidazole, pH 8) and the 6His-SUMO tag removed by incubation with the SUMO protease (1:100 ratio) 50–150 μg/ml followed by affinity chromatography [14]. Cleaved products were further purified by S-75 size exclusion chromatography (SEC buffer-50 mM sodium phosphate, 50 mM sodium chloride, 1 mM DTT, pH 8) (GE Healthcare Life Sciences). The purity of the protein was inspected using a 15% SDS PAGE (**Figure S1**). Isotope labeling of NHR2 was achieved in ~98% D2O-containing M9 media supplemented with^13^C-glucose and^15^N-NH_4_Cl as sole carbon and nitrogen sources, respectively. The labeled protein was purified with the same protocol as the unlabeled one. The M5 variant was purified with the same protocol as the NHR2 domain except for the SEC step: After SUMO cleavage, the M5 variant was precipitating; thus, high salt was used for the SEC step (SEC buffer-50 mM sodium phosphate, 300 mM sodium chloride, 1 mM DTT, pH 8). The concentration of NHR2 was determined (at 280 nm) using the molar extinction coefficient of 19480 M^−1^ cm^−1^ with molecular weight of 8547 Da, and 2980 M^−1^ cm^−1^ and 8100 Da for the M5 variant, respectively, calculated using the online server ProtParam.

### Inhibitor compound 7.44

The compound **7.44 (Figure 1D)** was kindly provided by the Developmental Therapeutics program, National Cancer Institute, USA (NCI code 162496). The **7.44** molecule was dissolved in DMSO-d6 as a 100 or 150 mM stock solution and stored at −20 °C for future use. The purity of the compound was shown to be >90% by LC-MS (**Figure S2)** and NMR techniques (**Figure 4A)**, and it was used without further purification.

### Size exclusion chromatography

Size exclusion chromatography (SEC) was performed with a Superdex 75 Increase 10/300 GL (GE Healthcare Life Sciences) column connected to ÄKTApure (Cytiva Life Sciences). Column efficiency was tested as described by the company [15]. The sample was prepared with 30 μM of NHR2 alone and in complex with a 1:1, 1:2, and 1:5 molar stoichiometric ratio of **7.44**, prepared in 50 mM sodium phosphate, 300 mM sodium chloride, 0.5 mM tris(2-carboxyethyl)phosphine, 3% (v/v) DMSO, pH 7.5 with a total volume of 250 μl, and filled into a 1 ml sample loop operated at a flow rate of 0.7 ml/min. The column was equilibrated with the same buffer as the sample buffer. Samples were incubated at 37 °C for 2 hours and then incubated overnight at room temperature before the experiment.

### Differential scanning fluorimetry

The method was applied as described in Kroeger *et al.*[16]. In short, the fluorescent dye SYPRO Orange interacts with hydrophobic surfaces of denatured proteins. This interaction changes the quantum yield of the dye [17, 18]. The fluorescence was measured with the real-time thermocycler qTOWER 2.0 (Analytik Jena AG, Germany). Here, 0.2 mg/ml NHR2 containing a His-tag were mixed with SYPRO Orange (1:1000) and **7.44** with concentrations ranging from 143 μM to 5.72 mM. The fluorescence signal was measured at the starting temperature (25 °C). The temperature was then increased by 1 °C/min up to 100 °C. With every increase in temperature, the fluorescence was measured. Every measurement was performed at least eight times independently.

### Circular dichroism spectroscopy

Far-UV circular dichroism (CD) spectra, used to investigate the thermal unfolding of NHR2 and the NHR2-**7.44** complex, were obtained using a JASCO Model J-815 CD spectrometer (Jasco, Japan) equipped with a temperature-controlled cell holder using a quartz cell with a 1-mm path length. The sample of 12.5 μM NHR2 for the apo form and, in addition, 200 μM of **7.44** for the complex was diluted in 10 mM sodium phosphate, 10 mM sodium chloride, 0.5 mM tris(2-carboxyethyl)phosphine, <1% (v/v) DMSO pH 6.5. The spectra were collected from the wavelength of 190–250 nm for apo NHR2 and 213-250 nm for the complex sample with accumulations of 20 scans at a speed of 200 nm min^−1^ at 25 °C. The thermal denaturation was recorded at the spectral minima of 222 nm by increasing the temperature from 25 °C to 95 °C in 1 °C increments with a slope of 1 °Cmin^−1^. CD curves were plotted using the Origin software (OriginLab Corporation, USA).

### Microscale thermophoresis

Prior to the MST experiments, purified NHR2 protein was labeled either with the dye Alexa Fluor^®^ 488 (NHS Ester Protein Labeling kit, ThermoFisher Scientific, USA) or Monolith NT™ (Protein Labeling Kit Blue-NHS, NanoTemper GmbH, Germany). The labeling was achieved by incubating 115 μM of protein and 350 μM of dye (465 μl reaction volume) for 1 h at room temperature, followed by 35 min at 30 °C in a buffer consisting of 20 mM sodium phosphate, 50 mM sodium chloride, pH 8.0 (MST buffer). The dye-labeled protein was purified using a PD-10 column containing Sephadex G-25 Medium (GE Healthcare Life Sciences) and then centrifuged at 50000 rpm for 30 min. The concentrations and the efficiency of the labeling were determined as indicated in the manual.

To determine the constant of dissociation of the NHR2 tetramer into dimers (*K*_tet_, **Eq. S1a**), 10 μl of 100 nM labeled NHR2 in MST-buffer (included with the capillary set from NanoTemper) was mixed with 10 μl unlabeled NHR2 with concentrations ranging from 10.3 nM to 337.5 μM, and the mixtures were incubated in the dark for 30 minutes.

To determine the *EC*_50_ of **7.44** binding to NHR2 protein, 100 nM of labeled NHR2 was mixed with **7.44** to concentrations ranging from 122 nM to 4 mM to the final volume of 50 μl (serial 1:1 dilution) in MST buffer containing 10% (v/v) DMSO. The samples were incubated overnight in the dark before the experiment. Each dilution was filled into Monolith NT.115 standard treated capillaries (NanoTemper Technologies GmbH). Thermophoresis was measured using a Monolith NT.115 instrument, with an excitation power of 50% for 30 s and MST power of 40% at an ambient temperature of 24 °C. Triplicates of the same dilution were measured. Microscale thermophoresis results were analyzed by the MO affinity analysis software (NanoTemper Technologies GmbH)

The data was analyzed and the *K*_tet_ (**Eq. 1**) or *EC*_50_ (**Eq. 2**) calculated using the MO affinity analysis software (NanoTemper, Germany), considering a 1:1 model.

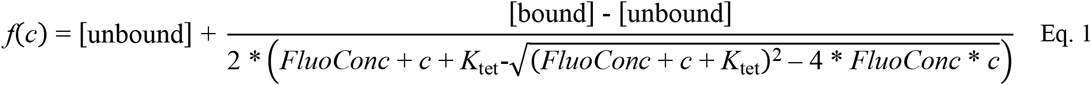

Here, *f*(*c*) is the fluorescence intensity, [*unbound*] is the concentration of unbound labeled and unlabeled NHR2, [*bound*] is the concentration of bound labeled and unlabeled NHR2, *FluoConc* is the concentration of labeled NHR2, *c* is the concentration of unlabeled NHR2.

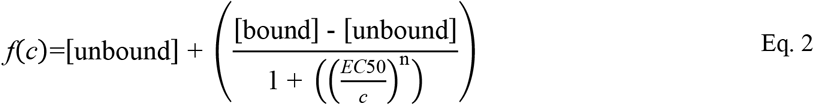

Here, *f*(*c*) is the fluorescence intensity, [*unbound*] is the concentration of unbound labeled NHR2, [*bound*] is the concentration of bound labeled NHR2, *c* is the concentration of **7.44**, *n* is the Hill coefficient. For **7.44** binding to NHR2, an additional 1:2 model included in the PALMIST software was considered to assess the MST data [19, 20]; this model takes into account the symmetry of an NHR2 dimer, which would allow binding of two **7.44** per dimer. The dissociation constant of **7.44** from an NHR2 dimer, *K*_lig_, was finally calculated using **Eqs. S1 to S12** by solving the equation systems with SymPy [21] and estimating the error with functions from Hans Dembinski [22].

### NMR spectroscopy

NMR experiments were performed at 298 K or 308 K on Bruker Avance III HD spectrometers operating at 600 MHz, 700 MHz, or 750 MHz, equipped with 5 mm triple resonance TCI (^1^H, ^13^C, ^15^N) cryoprobes and shielded z-gradients. Data was processed on NMRPipe [23] and analyzed by NMRView [24] software or TopSpin 3.2 (Bruker BioSpin). Sodium 2,2-dimethyl-2-silapentane-5-sulfonate (DSS) was used for chemical shift referencing.

### Assignment of ligand 7.44

For the assignment of the ^1^H resonances of ligand **7.44,** we prepared a solution of 2 mM of the ligand in 20 mM sodium phosphate buffer, pH 7.4, containing 100 mM sodium chloride and 10% (v/v) D_2_O at 298 K. The assignment of the protons of the ligand was achieved by the analysis of the 1D ^1^H and 2D ^1^H,^1^H-TOCSY spectra and aided by ^1^H chemical shift prediction as performed by the ChemNMR package, contained within the software ChemDraw (PerkinElmer). The 2D ^1^H,^1^H-TOCSY spectrum was acquired with 8 transients in a matrix with 2k data points in F2 and 256 increments in F1 with a relaxation delay of 1.0 s. “The obtained chemical shifts of ligand **7.44** were as follows: δH2”= 6.09 ppm (*s*); δH4”= 7.41 ppm (*J* = 1.7 Hz, d); δH3a= 3.66 ppm (*J* = 8.4; 17.5 Hz, dd); δH3b= 3.36 ppm (*J* = 7.3; 17.5 Hz, dd); δH2= 3.96 ppm (*J* = 7.7 Hz, t); δH4’= 6.89 ppm (*J* = 1.4 Hz, d); δH2’= 5.95 ppm (*s*); δH6’ and 7’= 6.86 – 6.81 ppm (*overlapped*); δH6”= 7.65 ppm (*J* = 6.8; 10.1 Hz, dd); δH7”= 6.98 ppm (*J* = 8.4 Hz, d).

### pKa value determination of 7.44

For the pKa value determination, 23 samples of 200 μM concentration of **7.44** were prepared from the pH range of 2 to 13 (pH of 0.5 steps) in 50 mM sodium phosphate, 100 mM sodium chloride, 10% (v/v) DMSO-d_6_. 1D ^1^H-NMR experiments were performed using 256 scans for each sample. ^1^H chemical shift values were extracted for the reporter protons using TopSpin 3.2 (Bruker BioSpin). The pKa was calculated using the chemical shifts of the protonated and unprotonated forms and their corresponding mole fraction values applying the Henderson-Hasselbalch equation as explained by Gift *et al.* [25]. The data were processed and plotted with the Origin software (OriginLab Corporation, USA).

### Saturation transfer difference NMR

For the STD-NMR experiments, two samples were prepared in 20 mM sodium phosphate, 50 mM sodium chloride, 0.5 mM tris(2-carboxyethyl)phosphine, 5% (v/v) DMSO_d6_, pH 6.5, and 10% (v/v) D_2_O, in which the ^13^C, ^15^N-NHR2 concentration was 50 μM and the ligand concentration was either 0 or 5000 μM for the reference and mixture, respectively. The STD-NMR spectra were acquired at 308 K with 512 scans. Selective saturation of protein resonances (on-resonance spectrum) was performed by irradiating at 0.6 ppm for a total saturation time of 2 s. For the reference spectrum (off-resonance), the samples were irradiated at −30 ppm. To obtain an STD effect, the subtraction was performed internally via phase cycling after every scan to minimize artifacts arising from temperature and magnet instability [26]. Proper control experiments were performed with the reference sample to optimize the frequency for protein saturation and off-resonance irradiation, to ensure that the ligand signals were not affected.

The STD effect was calculated using (*I*_0_ − *I*_STD_)/*I*_0_, in which (*I*_0_ − *I*_STD_) is the peak intensity in the STD spectrum and *I*_0_ is the peak intensity in the off-resonance spectrum. The STD intensity of the largest STD effect was set to 100% as a reference, and the relative intensities were determined [27]. The binding epitope is created by the comparison of the STD intensity relative to the reference one, and this relative intensity value is described by the STD amplification factor (A_STD_).

### Assignment of uniformly 2H, 13C, 15N-labeled NHR2 and interaction with 7.44

For 3D NMR experiments, 325 μM of uniformly ^2^H, ^13^C, ^15^N-labelled NHR2 in NMR buffer (20 mM sodium phosphate, 50 mM sodium chloride, 0.5 mM tris(2-carboxyethyl)phosphine, 10% (v/v) D_2_O, pH 6.5) was transferred into a 5 mm Shigemi tube (Sigma-Aldrich), and the experiments were performed at 308 K. Sequence-specific assignments were carried out using standard TROSY-based triple resonance experiments (HNCO, HN(CO)CA, HNCACB, HN(CO)CACB, ^1^H,^15^N-NOESY-HSQC, ^1^H,^15^N-TROSY-HSQC) [28, 29].

For the interaction experiments, two NMR samples, 47 μM of ^2^H, ^13^C, ^15^N-labelled NHR2 alone or in complex with 2 mM of **7.44**, were prepared in NMR buffer, which in addition contained 10% (v/v) DMSO_d6_, with a total volume of 200 μl and transferred into 3 mm NMR tubes.^1^H-^15^N-TROSY-HSQC experiment was performed using 216 scans per increment (about 19 hours) for both the samples at 353 K [28, 29].

### *In vitro* pharmacokinetics and toxicology prediction

The ADME property data was generated by Pharmacelsus GmbH, Saarbrücken, Germany. The methods of the six assays, aqueous solubility, plasma protein binding, plasma stability, hepatocyte clearance measurements, chemical stability, and Cytochrome P450 (CYP) inhibition, are described in detail in the Supporting Information. Further pharmacokinetic parameters were computed using the QikProp-Schrödinger suite (Release 2021-2) [30]. A computational toxicology assessment was performed using the toxicology modeling tools DEREK Nexus^®^ (Derek Nexus: 6.0.1, Nexus: 2.2.2) [31] and SARAH Nexus^®^ (Sarah Model - 2.0 Sarah Nexus: 3.0) [32] from Lhasa Limited, Leeds, UK.

### Molecular dynamics simulations

To investigate interactions between **7.44** and NHR2 in full atomic detail, we performed unbiased molecular dynamics (MD) simulations of **7.44** binding to NHR2. We used the tetrameric crystal structure of NHR2 (PDB ID 1WQ6); hereafter, we follow the residue numbering of the whole fusion protein RUNX1/ETO [1]. The protein structure was prepared using Maestro [33] for pH 7.4 to mimic physiological conditions. All titratable residues are present in the ionic state, except for histidines, which were assigned to the HID state. The initial 3D structure of **7.44** was prepared according to the 2D structural formula in **Figure 1D** and using the *S*-configuration. The initial 3D structure was geometry optimized at the HF/6-31G* level of theory using Gaussian 09 [34]. The optimized 3D structure was used for subsequent binding simulations. To do so, we randomly placed the tetrameric NHR2 structure, two **7.44** molecules, and sodium ions using PACKMOL [35]. The systems were solvated with the TIP3P solvent model, also using PACKMOL [35]. All relevant system files for subsequent simulations were generated using LEaP of Amber17 [36]. That way, we prepared a set of 35 independent initial configurations for MD simulations of **7.44** binding to NHR2. We applied the Amber ff14SB force field [37] for the protein and the GAFF force field [38] for **7.44**. Missing atomic partial charges of **7.44** were derived according to the restraint electrostatic potential (RESP) fit procedure [39, 40]. Ion parameters were taken from reference [41].

Details of the minimization, thermalization, and equilibration protocol are reported in ref. [42], which was successfully applied to study ligand binding processes [43, 44]. In short, the solvated systems were subjected to three rounds of energy minimization to eliminate bad contacts. Subsequently, systems were heated to 300 K, and the pressure was adapted such that a density of 1 g cm^−3^ was obtained. During thermalization and density adaptation, we kept the solute fixed by positional restraints of 1 kcal mol^−1^ Å^−2^, which were gradually removed. Subsequently, the systems were subjected to unbiased production simulations of 1000 ns length each to study **7.44** binding to NHR2 at a **7.44** concentration of ~1.2 mM. During these simulations, all molecules were allowed to diffuse freely without any artificial guiding force. All minimization, equilibration, and production simulations were performed with the pmemd.cuda module [45, 46] of Amber17 [36]. During production simulations, we set the time step for the integration of Newton’s equations of motion to 4 fs following the hydrogen mass repartitioning strategy [47]. Coordinates were stored into a trajectory file every 200 ps.

To identify epitopes on the NHR2 tetramer, where **7.44** preferentially binds, we computed occurrence propensities of heavy atoms of **7.44** around the tetramer structure (using the grid command in cpptraj and a grid spacing of 0.5 Å). For this, only frames were considered where any heavy atom of **7.44** is within 4 Å of any heavy atom of NHR2.

To observe how **7.44** interacts with the NHR2 tetramer, we first recorded for the whole trajectory, with which residues the ligand interacts (distance between heavy atoms < 4 Å). This constitutes for each frame a 244 bits vector. For further analysis, we only considered frames where the ligand interacts with at least five residues. After accounting for the internal symmetry of NHR2 by orienting all frames to a standard orientation, we applied a hierarchical clustering approach as implemented in scipy, using the Jaccard distance as a metric and a distance cutoff between cluster centroids of 0.3. The three most populated clusters from this approach correspond to the areas of highest propensity shown in **Figure 5**.

Details about the setup and performance of MD simulations for calculating the PAMPA permeability of **7.44** in its protonated and deprotonated form can be found in the Supplemental Notes.

## Results

### Expression, purification, and oligomer state of the NHR2 domain and the M5 variant

The NHR2 domain was overexpressed and purified, and the resulting material yielded homogeneous, monodisperse species in solution. The sample was homogenous as analyzed by size exclusion chromatography (SEC) (**Figure 2A**), SDS-PAGE (**Figure S1)**, and dynamic light scattering (DLS) (**Figure S3**). The oligomeric state of purified NHR2 was assessed by analytical ultracentrifugation (AUC) (**Figure S4**). The sedimentation profiles of NHR2 show a single major species with a sedimentation coefficient (*S* value) of 3.07, which corresponds to the molecular weight of 37.3 kDa. This value is in agreement with the calculated molecular weight of an NHR2 tetramer of 34.2 kDa. The weight percent contribution of the tetramer in 35 μM of NHR2 is 94% as obtained by integration of the continuous sedimentation coefficient distribution function. To check the function of the purified NHR2, we performed isothermal titration calorimetry (ITC) with the HEB fragment of E proteins (HEB (176-200)). The NHR2 domain has been shown to interact with the HEB fragment [48]. The resulting isotherm (**Figure S5**) yielded a binding enthalpy Δ*H* = −15 ± 6 kcal/mol and a dissociation constant *K*_*D*_ = 30 ± 4 μM, and this *K*_*D*_ value is in the same range as previously reported [48].

**Figure 2.**
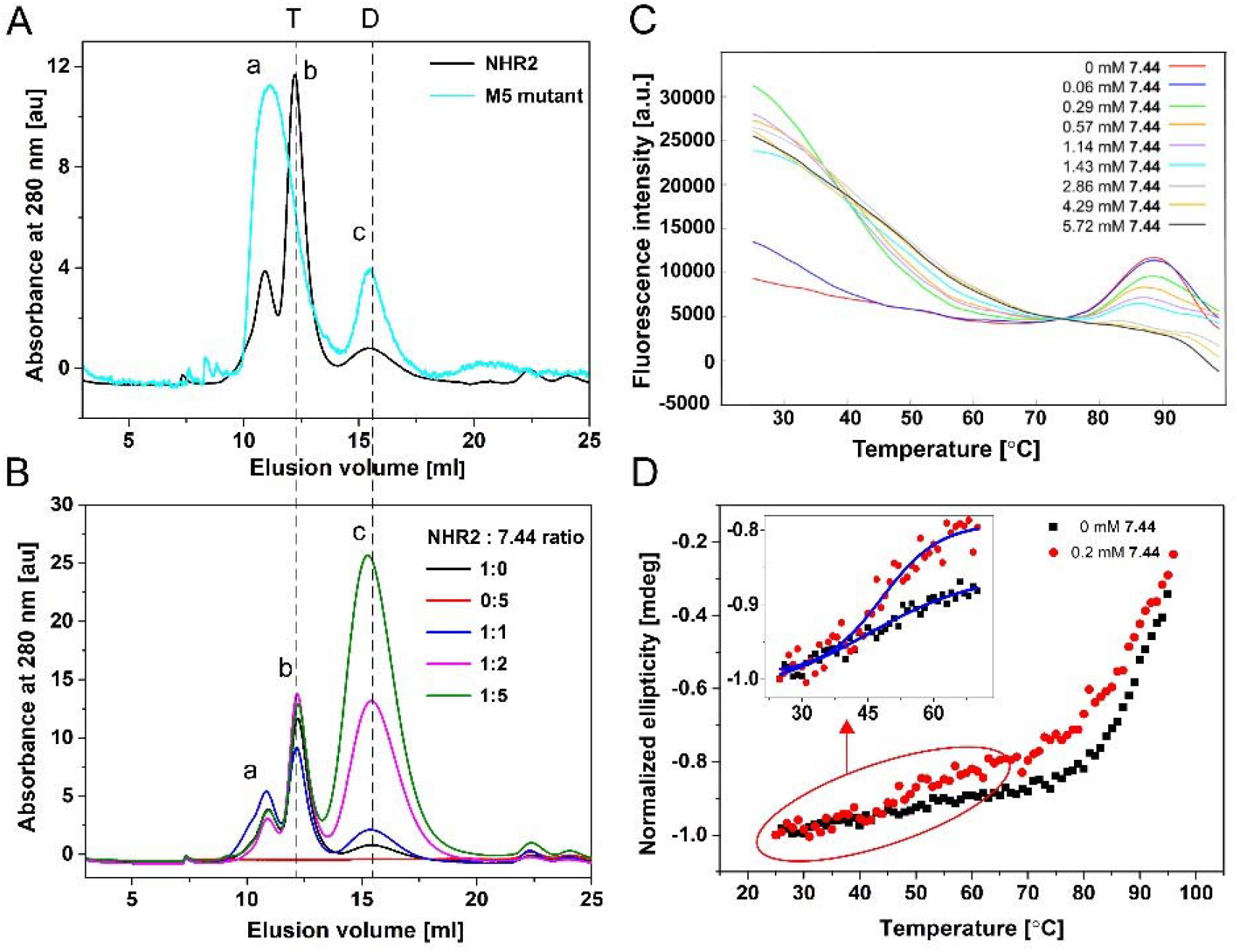
Stability of the NHR2 tetramer with and without 7.44 studied by size exclusion chromatography, DSF, and CD spectroscopy. **A)** Elution profiles of purified proteins of the NHR2 domain (black) and M5 variant (cyan) from size exclusion chromatography (SEC) obtained at 280 nm absorbance: Higher-order oligomers (a), tetramer (b), and dimer (c). The integration of peaks (b) and (c) of the NHR2 domain yields 92% of the tetramer and 8% of the dimer. **B)** Elution profiles of 30 μM of apo NHR2 (black), 150 μM of **7.44**, and NHR2:**7.44** with molar ratios of 1:1 (blue), 1:2 (magenta), and 1:5 (green) obtained from SEC at 280 nm. Samples were incubated at 37 °C for 2 hours and then incubated overnight at room temperature prior to the experiment. **C)** Fluorescence intensity in the presence of **7.44** at different concentrations; for better visualization, the curves were shifted along the y-axis to match them at the point between the two melting points of dimer and tetramer. **D)** Thermal denaturation of apo NHR2 (black) and NHR2 in the presence of **7.44** (red) monitored by CD spectroscopy at 222 nm wavelength. The CD data was normalized with respect to the highest ellipticity value for better comparison.

The SEC profile of NHR2 showed peaks that likely correspond to higher-order oligomer (a), tetramer (b), and dimer (c) (black in **Figure 2A**). Integration of peaks (b) and (c) reveals a relative abundance of 92% for the tetramer and 8% for the dimer, similar to the AUC data. These data are consistent with previously published sedimentation results [7, 10]. Earlier it was shown that the M5 variant forms a dimer but not a tetramer [10]. This finding is concordant with the crystal structure of NHR2 (PDB ID 1WQ6) [7], which reveals a four-helix bundle formed by four amphipathic α-helices, consisting of a symmetric dimer of dimers, where monomers in a dimer are oriented in a head-to-tail orientation. The M5 variant was overexpressed, purified, and characterized by SEC (**Figure 2A**), SDS-PAGE (**Figure S1**), and AUC (**Figure S4**). The SEC profile of the M5 variant showed higher-order oligomers (a), a potential contribution of tetramers (b), and dimer (c) (cyan in **Figure 2A**). The high intensity of elution peak (a) is concordant with the increased instability of the variant, which shows a higher tendency to form oligomers. AUC data of the M5 variant showed a slightly greater molecular weight compared to the theoretical one. Therefore, to confirm the dimeric state of the M5 variant, we performed AUC for the SUMO-M5 fusion protein, where we observed a measured molecular weight of 43.4 kDa, which agrees very well with the theoretical one of 43.9 kDa for the dimer (**Figure S4**).

### Influence of 7.44 on the NHR2 tetramer stability

Next, we analyzed the stability of the NHR2 tetramer in the presence of **7.44** by SEC, differential scanning fluorimetry (DSF), and CD spectroscopy [16, 49]. The SEC chromatograms indicate that increasing **7.44** concentrations lead to an increase in the relative ratio of NHR2 dimer to tetramer in the sample (**Figure 2B)**. **7.44** alone, measured as a control, does not show any absorbance (**Figure 2B**). Peak (c) of the dimer species increases with increasing **7.44** concentration, most probably due to a shift in the equilibrium from the tetramer to the dimer, indicating that **7.44** interferes with NHR2 tetramer stability. The decrease in the height of peak (b) of the tetramer species is not monotonic. Note, however, that the peak heights are inversely mirrored by those of peak (a) of higher-order oligomers. Due to the involvement of higher-order oligomers and the possibility that the absorption characteristics of **7.44** change when interacting with NHR2, which may contribute to the height of peak (c), we refrained from analyzing this data quantitatively.

In DSF, the thermally induced protein unfolding is monitored by the binding of the fluorescent dye SYPRO Orange [50] to the hydrophobic core of a protein that becomes exposed upon unfolding, leading to a related increase in fluorescence emission. The temperature at the midpoint of the unfolding transition is defined as the melting temperature (*T*_m_) of the protein [47]. Usually, interference of a ligand with a protein results in a shift in *T*_m_ of the protein compared to the native state [49] and monophasic melting [51]. However, we previously showed that the melting points of the NHR2 dimer and NHR2 tetramer are ~55 °C and ~85 °C, respectively [10]. Inhibition of NHR2 tetramer formation by **7.44** should alter the proportions of the two species in a dose-dependent manner towards the dimer [11], as also suggested by the SEC results. In such cases, rather than a monophasic melting at an intermediate melting temperature, a biphasic melting can occur, with NHR2 appearing to melt at the melting temperatures of the dimer and the tetramer [51]. Over a concentration range of **7.44** of a factor of ~100, melting occurs at 84-85 °C (**Figure 2C, Figure S7**), suggesting that this melting point relates to the tetramer species. The height of the fluorescence peak associated with this melting decreases with increasing **7.44** concentration (**Figure 2C**), indicating that the NHR2 tetramer concentration in the equilibrium declines. As to the melting of the dimer species at ~55 °C, an increase in the fluorescence signal at that temperature with increasing **7.44** concentration is observed, indicating that the NHR2 dimer concentration in the equilibrium rises. However, no clear melting points can be detected, because of an overlay with a strong and, with increasing temperature decreasing, fluorescence signal that has its maximum at ~30 °C. The fluorescence signal at that temperature also roughly increases with **7.44** concentration. This fluorescence signal may be explained in that dimeric NHR2 exposes its hydrophobic protein-protein interface [10] to which SYPRO Orange can bind; a temperature increase disfavors SYPRO Orange binding, which leads to a decrease in fluorescence [16]. Although these results demonstrate that **7.44** interferes with NHR2 tetramer stability and leads to a decline in NHR2 tetramer concentration with increasing **7.44** concentration, a quantitative analysis of this data is compromised due to the lack of a clear melting point at ~55 °C [51].

We also examined the thermal stability of NHR2 in the absence and presence of **7.44** by CD spectroscopy. The minima at negative ellipticity values at 208 nm and 222 nm wavelength indicate the presence of an α-helical structure in NHR2 in the absence of **7.44**(**Figure S6**), as expected from the crystal structure [7]. Upon addition of **7.44**, the ellipticity curve followed the same trend as for free NHR2, demonstrating that the α-helical structure is retained in the presence of **7.44** (**Figure S6)**; likewise, the dimeric M5 variant also retained the α-helical structure [10]. Note that due to the high voltage in the detector caused by the presence of DMSO in the buffer, the spectrum was recorded only from 213 nm to 250 nm wavelength. Therefore, the impact on the thermal stability of NHR2 by **7.44** was monitored at 222 nm wavelength (**Figure 2D**). Both *apo* NHR2 and NHR2 in the presence of **7.44** exhibited a cooperative biphasic melting (**Figure 2D**). The melting point (*T*_*m*_) of the first transition is ~48 °C, indicative of melting of the dimer, and the second *T*_*m*_ is ~88 °C, indicative of melting of the tetramer [10, 51]. The *T*_*m*_ of the tetramer is similar to the result from DSF (see above) and previous CD experiments [10]. For the dimeric M5 variant, a melting point of 60 °C was reported, associated with a broad transition between the folded and unfolded state [10]. The first phase transition is more pronounced in the presence of **7.44**, indicating a higher population of dimers due to the interference of **7.44** with NHR2. In addition, the second phase transition of NHR2 in the presence of **7.44** occurs at a lower temperature and is broader, suggesting a less tight association between the NHR2 a-helices.

To conclude, these results demonstrate that **7.44** interferes with NHR2 tetramer stability. Furthermore, the increase in the dimer population upon the addition of **7.44** indicates that **7.44** fosters a tetramer-to-dimer transition of NHR2.

### Dissociation constants of the NHR2 tetramer and of 7.44 with respect to the NHR2 dimer

For determining the dissociation constant of **7.44** binding to NHR2, *K*_lig_ (**Eq. S1b**), the tetramer-to-dimer equilibrium upstream of the ligand binding needs to be considered (**Eq. 4**), characterized by the dissociation constant *K*_tet_ (**Eq. S1a**). For that, we assume that tetrameric NHR2 (T) is in equilibrium with its dimer (D) and that **7.44** (L) binds to D after dissociation of T, leading to **Eq. 4**. Because of the large distance between the anticipated binding sites at the hot spot regions (**Figure 1C**), we furthermore assume that *K*_lig_ of the first and second ligand binding events are identical.

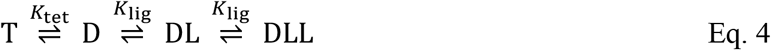

We determined *K*_lig_ by microscale thermophoresis (MST) measurements as a function of *K*_tet_ and *EC*_50_ of **7.44** binding to NHR2 according to equations S1-S12 [52, 53]. MST is based on measuring the directed movements of molecules along a generated temperature difference (thermophoresis). The movements depend on the combined impact of size, charge, and solvation shell of the molecules. For determining *K*_tet_ and *EC*_50_, a constant concentration of fluorescently labeled NHR2 was measured in the presence of varying concentrations of the titrants, unlabeled NHR2 or **7.44**, respectively. With increasing concentration of the titrant, the fraction of labeled NHR2 that forms a complex with the titrant will increase.

To determine *K*_tet_, unlabeled NHR2 was titrated to 50 nM of labeled NHR2 in 16 concentrations ranging from 10.3 nM to 337.5 μM. MST was recorded for each sample, and a non-linear regression curve was fit with the *K*_D_ Fit equation (**Eq. 1**) (**Figure 3A**). The inflection point of the curve revealed *K*_tet_ = 11.3 ± 1.81 μM. Next, we titrated 16 concentrations ranging from 122 nM to 4 mM of **7.44** to 100 nM of labeled NHR2. Based on the calculated *K*_tet_, more than 99% of the protein is in a dimeric state at this NHR2 concentration. This calculation was confirmed by a microfluidic diffusional sizing experiment, where 100 nM of dye-labeled NHR2 showed only dimer size, according to the measured hydrodynamic radius of 2.22 nm, with a calculated molecular weight of 16.74 kDa (**Figure S8**). After overnight incubation of the samples, MST was recorded for each sample (**Figure S9**). To obtain *K*_lig_, the MST data was analyzed in two independent ways. First, the resulting binding data was fit to a non-linear regression curve to obtain the *EC*_50_ value (**Eq. 2**) (**Figure 3B**). The inflection point of the curve revealed an *EC*_50_ of 2.5 ± 0.8 μM (**Table 1**), demonstrating binding of **7.44** to NHR2 and suggesting a slow dissociation kinetic [19]. The range of Hill coefficients *n* = 1.0-1.5 obtained from the fits reveals no or only weakly cooperative binding of **7.44** to NHR2. A control experiment was performed using the same concentrations of **7.44** together with the labeling dye, but without NHR2. A *K*_lig_ = 3.95 ± 1.28 μM was subsequently computed using **Eqs. S11 and S12**. A simulation of the species distribution using *K*_lig_ and *K*_tet_ shows how both constants result in a *EC*_50_ of 2.5 μM, which was used as a parameter to calculate *K*_*lig*_ (**Figure S11**). As the majority of NHR2 is in a dimeric state during the MST experiments and assuming that the MST signal is dominated by the formation of DL (**Eq. 4**), one can expect that *EC*_50_ ≈ *K*_lig_; these values are in the range of 2 μM to 4 μM and similar within experimental uncertainties. Second, as a further control, *K*_lig_ was obtained from MST data by considering a 1:2 binding model, assuming that two **7.44** can bind to the symmetric NHR2 dimer, as implemented in the PALMIST software [19, 20]. This yielded *K*_lig_ = 2 μM, 68.3% CI [0,6] μM **(Figure S10)**, in the same range as *K*_lig_ and *EC*_50_ above, although with high uncertainty (see **Figure S10** for details).

**Figure 3.**
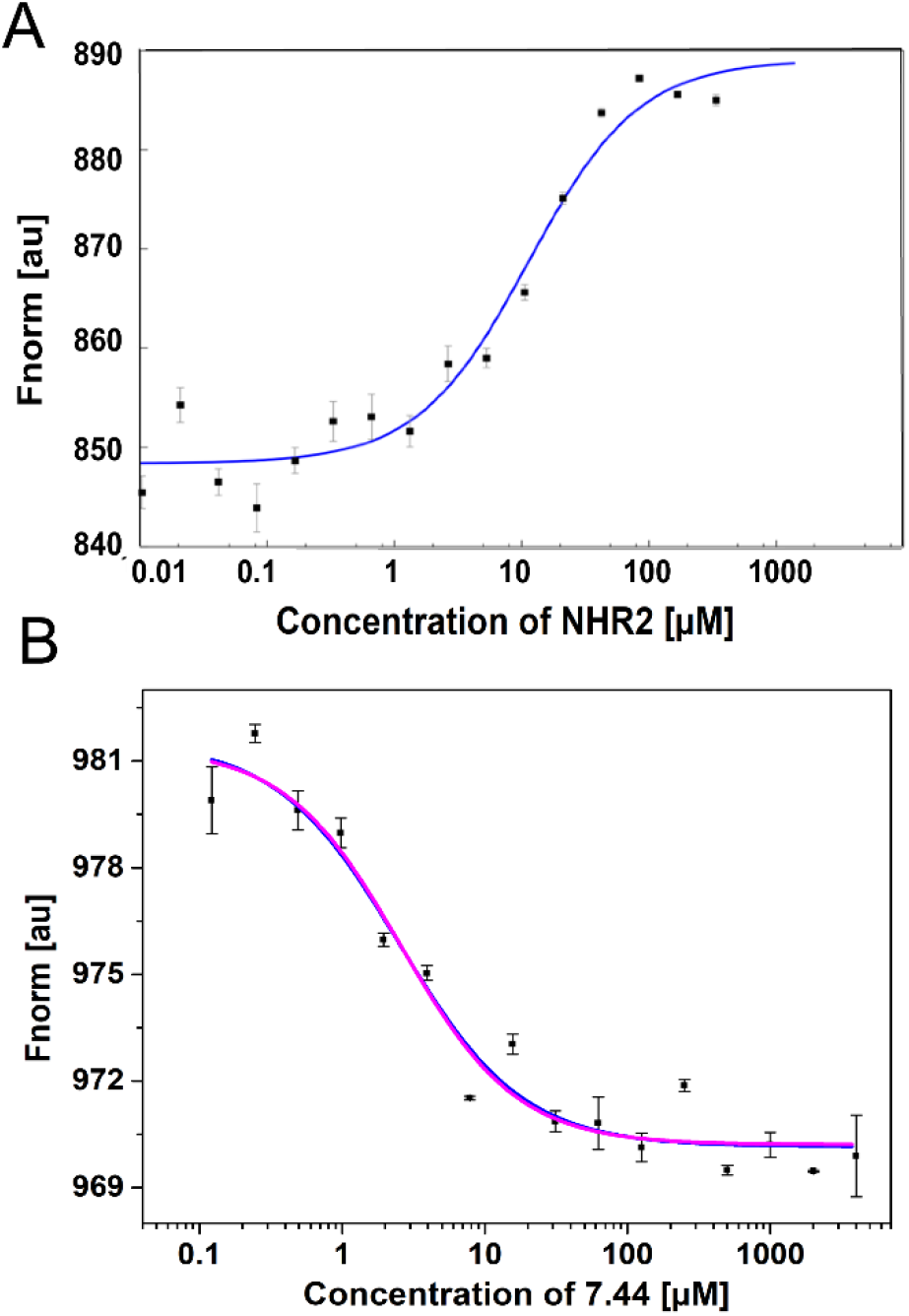
7.44 binding to NHR2 detected with MST. **A)** Titration of unlabeled NHR2 to a constant concentration of labeled NHR2 induces a change in thermophoresis. *K*_tet_ = 11.3 ± 1.81 μM was calculated using **Eq. 1**. **B)** Titration of **7.44** to a constant concentration of labeled NHR2 induces a change in thermophoresis. *EC*_50_ was calculated by fitting the data points to **Eq. 2**. The fitted values are shown in **Table 1**. The range of Hill coefficients *n* = 1-1.5 was obtained from the fits. Error bars denote the standard error in the mean and were calculated from technical triplicate.

**Table 1.** Binding characteristics of **7.44** binding to NHR2 from MST experiments.^[a]^

| Analysis method | $EC_{50}$<sup>[b]</sup> |
| --- | --- |
| Thermophoresis | $2.6 \pm 0.8$ |
| Thermophoresis with T-jump | $2.5 \pm 0.8$ |
| T-jump | $2.1 \pm 0.7$ |

<sup>[a]</sup> $EC_{50}$ was calculated using **Eq. 2**.
<sup>[b]</sup> In $\mu\text{M}$ .

To conclude, to our knowledge for the first time, the equilibrium constant for the dissociation of the NHR2 tetramer to NHR2 dimers and *K*_lig_ for **7.44** binding to the NHR2 dimer were determined.

### Epitope mapping of 7.44 via STD-NMR

In order to understand how **7.44** interacts with NHR2, we used saturation transfer difference-NMR (STD-NMR) [26, 27] to identify which atoms of **7.44** come close to the protein when the complex is formed (ligand epitope mapping). A typical STD-NMR experiment relies on the ligand exchange between the bound and free states. The difference between a spectrum without saturation of protein resonances (reference spectrum) and a saturated one yields the STD spectrum, in which it is possible to identify i) if the ligand binds to the protein (there are signals in the STD-NMR spectrum), ii) which protons of the ligand are involved in binding (only these are visible), and iii) which protons of the ligand are closer to the protein upon binding (by comparing the relative signal intensities) [26, 27].

The reference ^1^H spectrum of **7.44** and the corresponding STD-NMR spectrum, acquired in the presence of 100-fold molar excess of **7.44**, are shown in **Figure 4A** and **B**. All **7.44** protons show a signal in the STD-NMR spectrum, which is a clear indication that **7.44** binds to NHR2, in accordance with MST data. Different protons receive differential amounts of saturation (see color scale in **Figure 4B**), which allows mapping the epitope of the ligand. Proton H7′′ shows the highest saturation and was thus set to 100%, indicating its closest proximity to the protein. Other protons of the two ring systems show relative intensities between 64% and 97%, suggesting that **7.44** binds with both ring systems to NHR2. Protons H2 and H3, which are in the connecting alkyl chain, show the lowest detectable intensities of 48% and 20%, respectively. Protons more than 5 Å away from the protein normally do not appear in the STD-NMR spectrum.

To conclude, the STD-NMR data revealed that **7.44** interacts with NHR2 via both heteroaromatic moieties.

**Figure 4.**
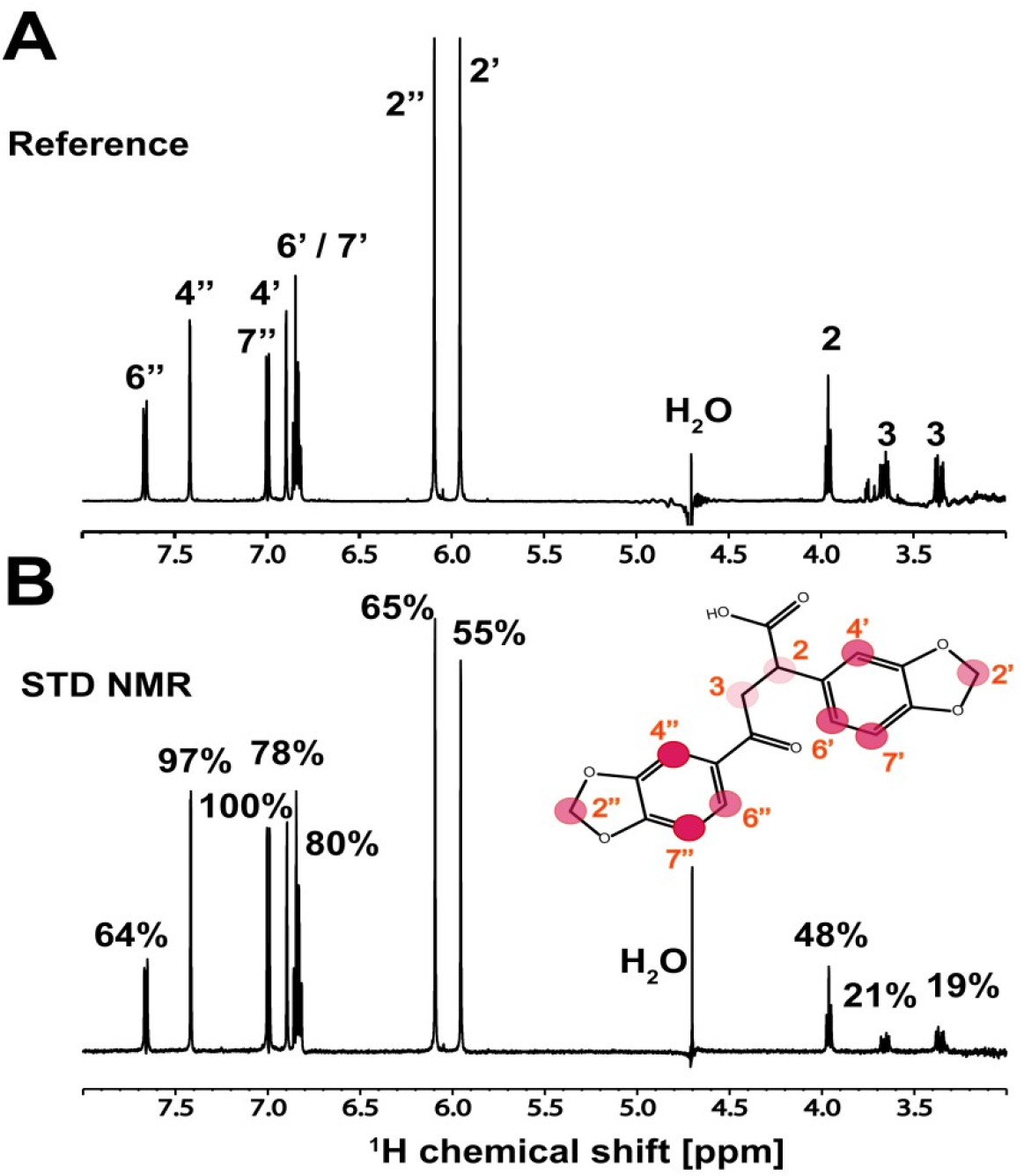
^1^H STD-NMR experiment of 7.44 binding to NHR2. **A)** Reference spectrum of **7.44**. Assignment of the individual peaks is indicated (see **7.44** structure in the bottom panel). **B)** ^1^H STD-NMR spectrum of ligand **7.44** in the presence of NHR2. Relative intensities are indicated on top of the corresponding peaks and mapped onto the structure of **7.44**. The circles are colored by the relative intensity of the protons, which is correlated with the proximity to the protein: red – 100% (the closest), lighter red– 97% to 64%, fade red – 48% to 19%. The proton numbering is indicated by the red numbers. Data was acquired at 600 MHz and 298K, with a 100-fold molar excess of **7.44**.

### The presence of 7.44 induces CSPs in the NHR2 N-terminus including the predicted tryptophan hot spot residues

Next, we proceeded to study the interaction of **7.44** with NHR2 from the protein side. We started by assigning the backbone resonance peaks of NHR2. We used a 325 μM sample of uniformly [^2^H, ^13^C, ^15^N]-labeled NHR2 and followed a standard triple-resonance-based approach [28, 29]. The NHR2 concentration in these experiments was about 30-times higher than *K*_tet_, shifting the equilibrium to the tetramer side. The solution NMR experiments of the tetrameric NHR2 protein were challenging due to the size and shape of the molecule [54]. The four inter-twined α-helices of 68 amino acids each led to a rod-like shape of 102 Å length (**Figure 1B)**. This extended conformation results in a large rotational diffusion anisotropy, causing a poor signal-to-noise ratio and only subtle peaks to be detected in the triple resonance experiments. Furthermore, the chemical shift dispersion of the backbone amide protons is limited to values between 7.5 ppm and 8.5 ppm. Albeit 61 backbone amide peaks were observed in the ^1^H-^15^N TROSY-HSQC spectrum (**Figure S12**), only 32% of these could be assigned sequence-specifically due to the low dispersion and low sensitivity in more complex (3D) experiments. The complexity of performing NMR experiments on coiled-coil assemblies is reflected in the PDB database where no solution NMR structure of a tetramer has been deposited (until October 2021) with a monomer size > 65 amino acids [54]. The number of amide cross-correlations in the ^1^H-^15^N TROSY-HSQC spectrum of ^2^H, ^13^C, ^15^N-labeled NHR2 (**Figure 5A**) is consistent with a single conformation, indicating that NHR2 forms a symmetric homotetramer in solution. No additional amide peaks were found after three weeks of storing the NMR sample at 35 °C, indicating that no alternative oligomeric structure formed over this time. The N-terminal residues from Q485 to D495, T519, and the C-terminal residues from Y544 to K552 residues were unambiguously assigned (**Figure 5A**). These regions are most important for evaluating 7.44 binding according to our binding mode model [11].

**Figure 5.**
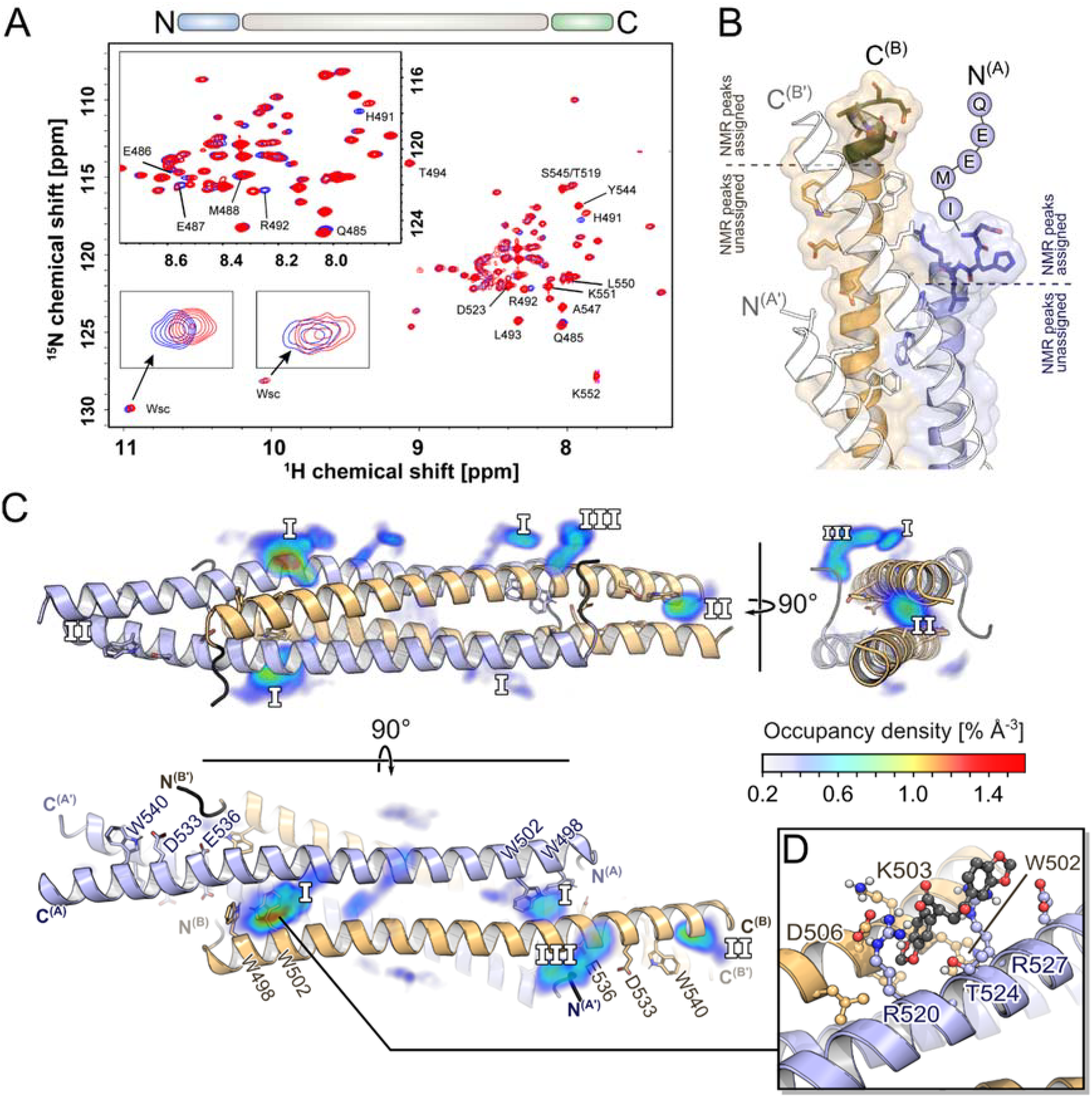
TROSY-HSQC NMR experiments of ^2^H, ^15^N, ^13^C-labeled NHR2 in the absence and presence of 7.44. **A)** 2D ^1^H-^15^N TROSY-HSQC spectra of 47 μM of apo NHR2 (blue) and in complex with 2 mM of **7.44** (red). Both spectra were recorded at 60 °C on a 750 MHz spectrometer for about 19 hours each and illustrate the changes in amide proton and nitrogen chemical shifts induced in NHR2 upon binding of **7.44**. 32% of amide signals from the N-(light blue; residues Q485 to D495, T519) and C-(light green; residues Y544 to K552) terminus were unambiguously assigned (depicted above the spectra); these residues are labeled with a single-letter code (see also panel B for the location). Note that the first five (QEEMI) and last four (DLKK) amino acids were also assigned but are not shown in the crystal structure. **B)** Chemical shift perturbations (CSPs) of residues Q485 to L493 (blue) upon addition of **7.44** mapped onto the NHR2 crystal structure (PDB ID 1WQ6). Residue R492, which shows the highest CSP, is associated with three hotspot residues (see text). **C)** Occurrence propensities of **7.44** interacting with the NHR2 tetramer from MD simulations; the densities are mapped around the NHR2 crystal structure (PDB ID 1WQ6). Assigned residues at the termini showing CSP are colored in black. The side chains of the previously identified hotspots are shown as sticks in the figure. The preferred binding epitopes with the highest densities are labeled as **I**, **II**, and **III**. Identical binding epitopes due to the internal symmetry of the NHR2 tetramer are labeled identically. Here, ligands binding to epitope **I** and **II** form interactions with the hotspots W502 and W540, respectively. **D)** Representative binding pose of **7.44** in epitope **I**. The 1,3-benzodioxole moiety inserts in between the helices, close to the hotspot residue W502. Identical binding poses are observed for all epitopes **I**.

To investigate the interaction of **7.44** with NHR2, NMR samples of ^2^H, ^13^C, ^15^N-labeled NHR2 (47 μM) in the absence and presence of **7.44** (2 mM) were prepared in 10 % (v/v) DMSO, and ^1^H-^15^N TROSY-HSQC spectra were measured at 60 °C for 19h. **Figure 5A** shows the superposition of the spectra. Considerable chemical shift perturbations (CSPs) occur for the N-terminal residues Q485 to L493 (colored in blue in **Figure 5B**); no CSPs were found for residues at the C-terminus. Unassigned peaks also showed CSPs. In addition, the protons at the indole nitrogen of tryptophan residues (Wsc) also displayed minor CSP (insert in **Figure 5A**).

The most affected residues are H491 and R492 (**Figure 5A, B, D)**. R492 is likely important for tetramerization as it forms a salt bridge with D533 and a solvent-exposed salt bridge with E536 that, in turn, interacts by a charge-assisted hydrogen bond with the indole NH of W540 [11]. Hence, the CSP suggest that **7.44** interferes with these interactions. Even though the backbone chemical shift assignments are also missing for the other two hotspot tryptophane residues (W498, W502; NHR2 contains three tryptophans in total), weak CSP of tryptophan side chains (Wsc) suggest that at least one (or even two) hotspot residue(s) is (are) involved in binding or experience a change in the environment (insert in **Figure 5A**). No signals for a third Wsc were observed, likely due to signal overlap or a low signal-to-noise ratio.

To conclude, CSP of NHR2 residues in the presence of **7.44** indicate interactions or conformational changes, particularly involving R492 in the N-termini, which is associated with three hotspot residues. At least one or two tryptophan side chains are involved; all three tryptophans of NHR2 were predicted as hot spots [11].

### Prediction of molecular interaction patterns of 7.44

To predict how **7.44** elicits its effect on NHR2 on a molecular level, we performed a series of MD simulations of free diffusion of **7.44** around the tetramer. We focused on the tetramer to mimic the situation in the NMR experiments where no NHR2 dimer was detectable. In these simulations, no biasing force was applied to any of the molecules. We observed multiple events of **7.44** binding to and unbinding from the NHR2 tetramer on visual inspection of the MD trajectories. By calculating occurrence propensities of **7.44** when the compound is in contact (distance between non-hydrogen atoms of **7.44** and NHR2 < 4 Å) with the NHR2 tetramer, distinct epitopes where **7.44** preferably forms contacts with the NHR2 tetramer emerge (**Figure 5B**). Due to the internal symmetry of the NHR2 tetramer, there are up to four identical locations for each epitope. Among these, epitope **I**, located between helices A and B of one dimer, is the most populated one. Here, **7.44** inserts its 1,3-benzodioxole moiety between the helices to form interactions with W502 (**Figure 5C**). In epitope **II**, only moderately less populated, the ligand intercalates between the helices A and A’ of the opposing dimers with its 1,3-benzodioxole moiety in the vicinity of W540. In epitope **III**, the ligand mostly forms interactions with the N-terminal part of NHR2, which was resolved as an unstructured loop in the crystal structure.

To conclude, the highly populated epitopes **I** and **II** suggest that at least one of the 1,3-benzodioxoles forms interactions with one of two buried tryptophan residues in the tetramer complex that had been identified as hot spots before [11].

### pKa value determination of 7.44 by NMR

**7.44** contains a single acidic group. To determine its pKa value, NMR spectra are collected for a pH range from 2 to 13 using 200 μM of **7.44**. As the pH of the solution is changed, the position of the proton peaks in the ^1^H NMR spectrum shift (**Figure 6A)**. Sodium 2,2-dimethyl-2-silapentane-5-sulfonate (DSS) was used for chemical shift referencing and calibrated to a zero ppm value in each case. Using the assignment of **7.44** (**Figure 4A**), we identified H2 and H3 as reporter protons, which are close to the acidic group and displayed pronounced chemical shift changes over the pH titration. Then the pKa was calculated by using the chemical shifts of the protonated and unprotonated forms at the respective pH and fitting to the Henderson-Hasselbalch equation [25] (**Figure 6B**). This yielded pKa = 3.99 ± 0.01 and 3.93 ± 0.01 of the carboxylic group as indicated by proton H2 and H3, respectively.

**Figure 6.**
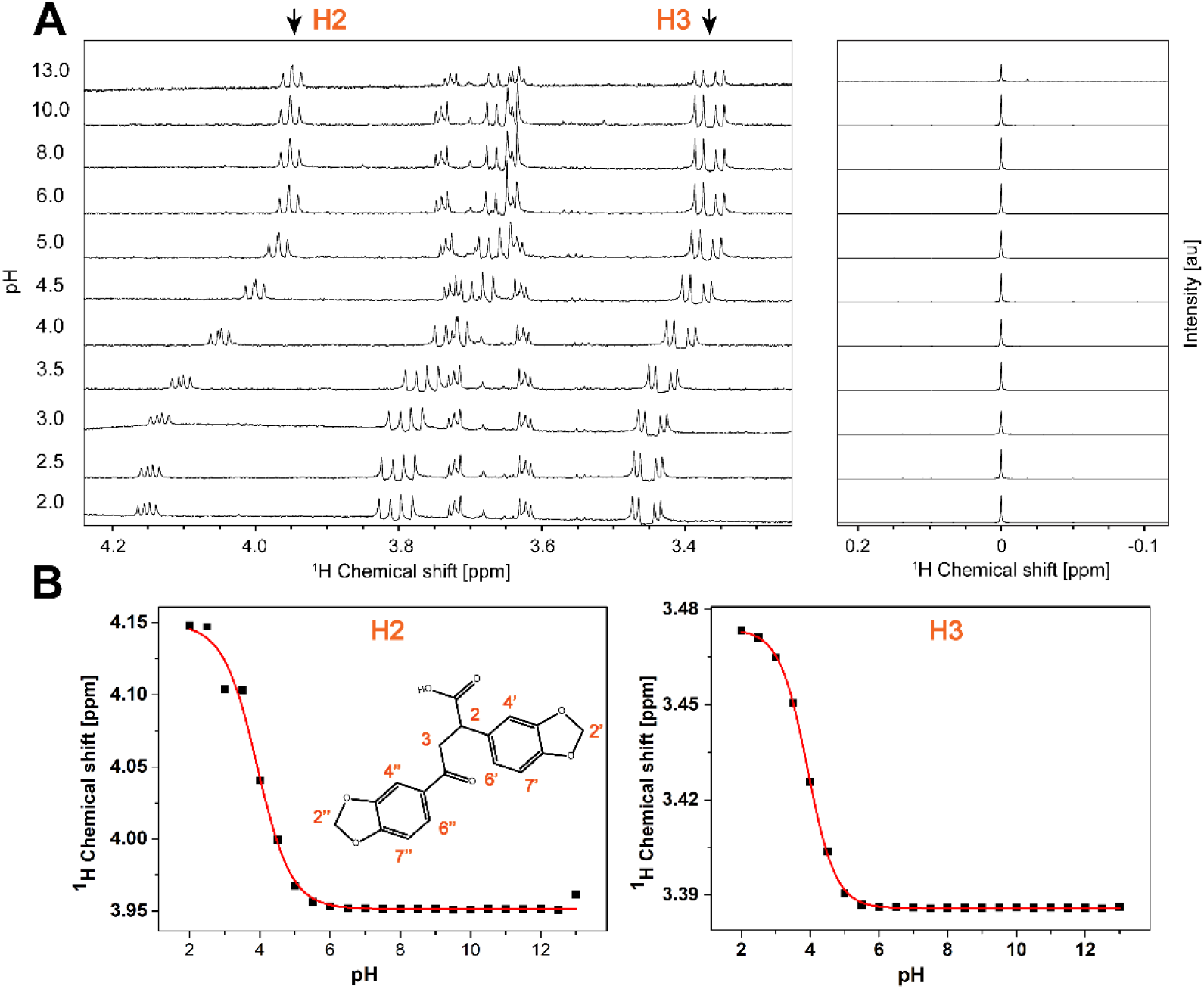
Determination of the pKa value for 7.44 by ^1^H NMR. **A)** ^1^H NMR spectra of 200 μM of **7.44** measured from pH 2 to 13 at 298 K at 600 MHz. Sodium 2,2-dimethyl-2-silapentane-5-sulfonate (DSS) was used for chemical shift referencing and calibrated to a zero ppm value. B) Titration curves obtained from the ^1^H chemical shifts of the protons H2 and H3 as a function of pH. The pKa values were calculated using the Henderson-Hasselbalch equation [25], yielding a pKa value of 3.99 ± 0.01 and 3.93 ± 0.01 of the carboxylic group as indicated by proton H2 and H3, respectively.

### Physicochemical, *in vitro* pharmacokinetic, and toxicological properties of 7.44

Next, we assessed physicochemical, *in vitro* pharmacokinetic, and toxicological properties of **7.44** (**Table 2, 3**). (1) *Kinetic solubility in PBS.* The aqueous solubility was determined in phosphate-buffered saline (PBS) containing 1% DMSO at pH 7.4 and yielded 177 ± 8 μM (60 μg/ml) for **7.44**. To achieve oral absorption, a compound with medium intestinal permeability and a projected human potency of 1 mg/kg needs a minimum aqueous solubility of 52 μg/ml [55]. (2) *Plasma protein binding.* Plasma protein binding was determined by the ultrafiltration procedure. This assay involves the separation of the free fraction from the bound fraction by a semi-permeable membrane with a high mass cutoff of 30 kDa. **7.44** showed a high (bound fraction: 98.4%) binding to plasma proteins from a mouse. Plasma protein binding influences the distribution of drugs in body tissues, and compounds with > 99% plasma protein binding are limited in terms of the amount of free compound that are available to act on the targeted tissue [56]. (3) *Plasma stability.* No degradation was observed for **7.44** when incubated in mouse plasma up to 4 h at 37 °C. Hence, plasma enzymes will not significantly alter the bioavailability of the compound. (4) *Hepatocyte stability.* Metabolic conversion of **7.44** was monitored by the disappearance of the compound in the presence and absence of hepatocytes from mouse. After 120 min of incubation time, 73.8% of **7.44** remained, demonstrating a half-life time > 120 min. An intrinsic clearance rate (C_Lint_) of 2.5 μl / min / million cells for **7.44** was obtained, indicating high hepatocyte stability. (5) *Chemical stability in PBS.* Chemical stability of **7.44** in PBS buffer at pH 7.4 was assessed for 4 h. No degradation was observed, suggesting that the compound is stable under physiological conditions for at least 4 h. (6) *Cytochrome P450 (CYP) inhibition.* Inhibition assays were performed for seven CYP isoforms at two different concentrations (1 μM and 10 μM, **Figure 7**). For **7.44**, there is a potential risk to inhibit CYP2B6, 2C9, 2C19, and 3A4 as indicated by a dose-dependent effect and inhibition > 50% at 10 μM. For the other tested enzymes CYP1A2, 2C8, and 2D6, the potential is considered low (inhibition < 25% at 10 μM resulting in estimated IC50 values of < 10 μM). (7) *Permeability prediction (PAMPA).* We predicted **7.44**’s permeability through a PAMPA membrane from molecular simulations and configurational free energy computations applying experimental data from nine reference compounds for calibration (**Figure 8A, Table S10**), as done before (see Supporting Information) [57]. The computed *P*_eff_ is 1.51 × 10^−6^ cm sec^−1^ for deprotonated (negatively charged) **7.44** and 8.51 × 10^−6^ cm sec^−1^ for the protonated (neutral) form (**Figure 8B**, **Table S11, Table 3**). This classifies **7.44** as low to medium (*P*_eff_ = 0.7 × 10^−6^ −2.1 × 10^−6^ cm sec^−1^) and highly permeable (*P*_eff_ > 4.7 × 10^−6^ cm sec^−1^) [58]. (8) *Prediction of further pharmacokinetic properties*. Using Qikprop from the Schrödinger software suite [30], we predicted the central nervous system activity, the brain-blood partition coefficient, and the IC50 value for blockage of HERG K^+^ channels (**Table 3**). The results classify **7.44** as almost central nervous system-inactive and predict no blockage of HERG K^+^ channels. Furthermore, no violations of Lipinski’s rule of five [59] or Jorgensen’s rule of three [60], for judging oral bioavailability, were found. Finally, no property of **7.44** computed by Qikprop falls outside the 95% range of similar values for known drugs. (9) *Toxicology prediction*. We predicted toxicological properties of **7.44** using the softwares DEREK Nexus [31] for a variety of endpoints and SARAH Nexus [32] for mutagenicity predictions (**Table 3**). No toxicophores were identified in **7.44**, and the compound is predicted to be inactive with respect to bacterial mutagenicity.

**Table 2.** Physicochemical and *in vitro* pharmacokinetic properties of **7.44**.

| No. | Description | Value |
| --- | --- | --- |
| 1 | Kinetic solubility (99% PBS, 1% DMSO) | $177 \pm 8 \mu\text{M}$ |
| 2 | Plasma protein binding (mouse plasma, 60 min) | $98.4 \pm 0.2\%$ |
| 3 | Plasma stability (mouse plasma, 0-240 min) | No degradation |
| 4 | Hepatocyte stability (mouse hepatocytes) | $2.5 \mu\text{l/min/million cells}$ |
| 5 | Chemical stability in PBS (0-4 h) | No degradation |
| 6 | Cytochrome P450 inhibition (at $1 \mu\text{M}$ and $10 \mu\text{M}$ concentration) | See Figure 7 |

**Table 3.**
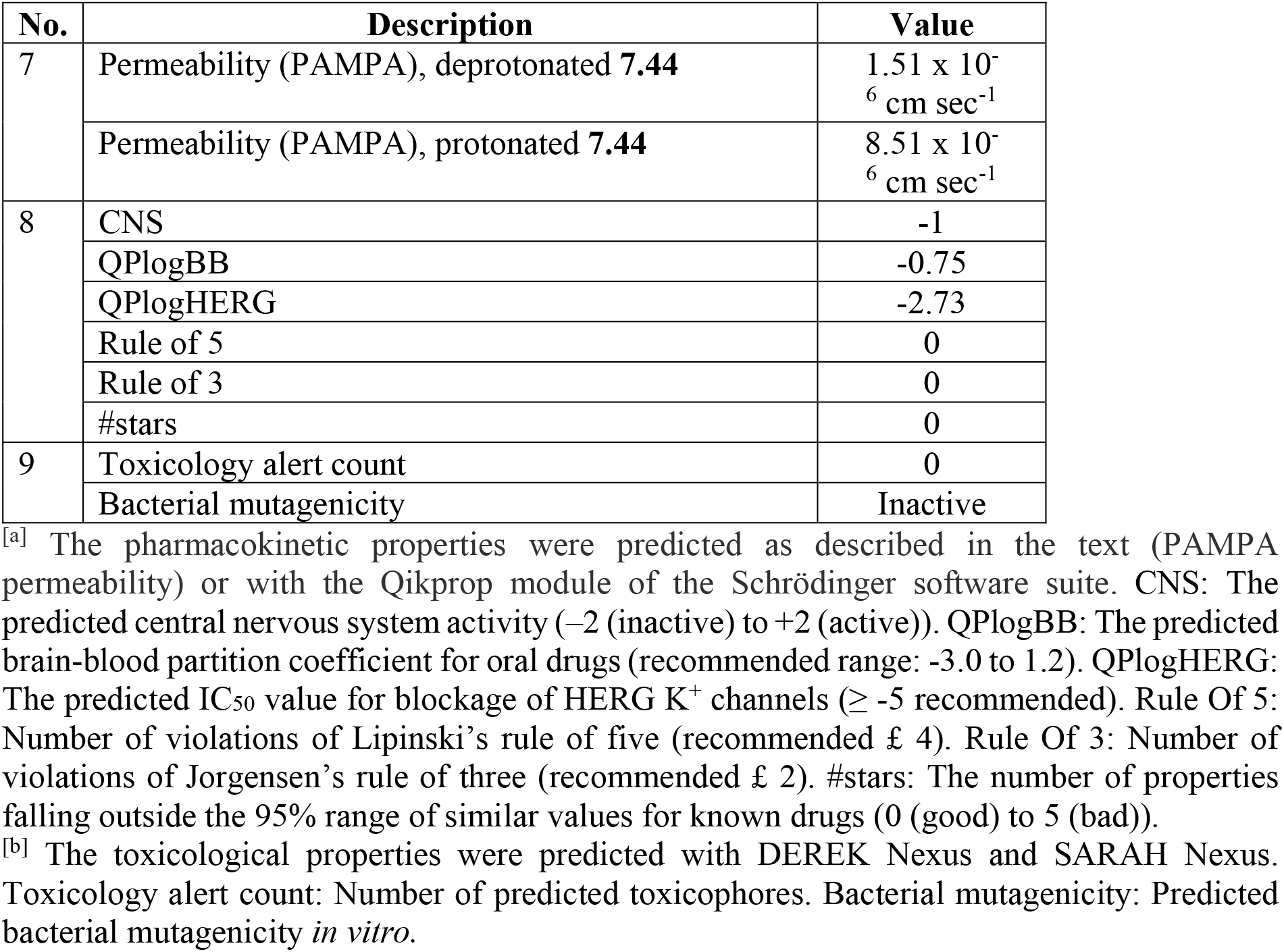
Predicted pharmakokinetic ^[a]^ and toxicological ^[b]^ properties of **7.44.**

| No. | Description | Value |
| --- | --- | --- |
| 7 | Permeability (PAMPA), deprotonated **7.44** | $1.51 \times 10^{-6} \text{ cm sec}^{-1}$ |
| | Permeability (PAMPA), protonated **7.44** | $8.51 \times 10^{-6} \text{ cm sec}^{-1}$ |
| 8 | CNS | -1 |
|  | QPlogBB | -0.75 |
|  | QPlogHERG | -2.73 |
|  | Rule of 5 | 0 |
|  | Rule of 3 | 0 |
|  | #stars | 0 |
| 9 | Toxicology alert count | 0 |
|  | Bacterial mutagenicity | Inactive |

<sup>[a]</sup> The pharmacokinetic properties were predicted as described in the text (PAMPA permeability) or with the Qikprop module of the Schrödinger software suite. CNS: The predicted central nervous system activity (−2 (inactive) to +2 (active)). QPlogBB: The predicted brain-blood partition coefficient for oral drugs (recommended range: −3.0 to 1.2). QPlogHERG: The predicted IC<sub>50</sub> value for blockage of HERG K<sup>+</sup> channels ( $\geq -5$ recommended). Rule Of 5: Number of violations of Lipinski's rule of five (recommended $\leq 4$ ). Rule Of 3: Number of violations of Jorgensen's rule of three (recommended $\leq 2$ ). #stars: The number of properties falling outside the 95% range of similar values for known drugs (0 (good) to 5 (bad)).
<sup>[b]</sup> The toxicological properties were predicted with DEREK Nexus and SARAH Nexus. Toxicology alert count: Number of predicted toxicophores. Bacterial mutagenicity: Predicted bacterial mutagenicity *in vitro*.

**Figure 7.**
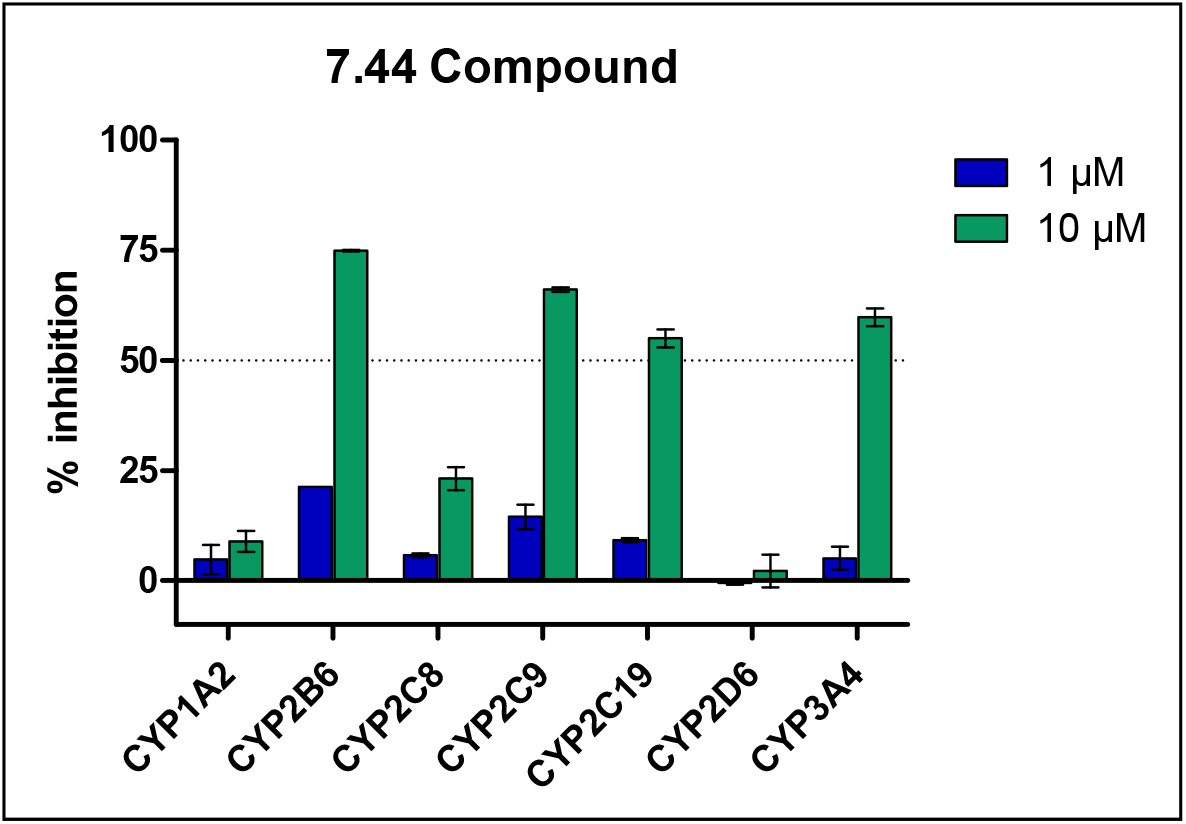
Cytochrome P450 (CYP) inhibition assay for 7.44. The inhibition assays were performed for seven CYP enzymes at two different concentrations, 1 μM (blue) and 10 μM (green). **7.44** showed a potential to inhibit CYP2B6, 2C9, 2C19, and 3A4. For the other tested enzymes, CYP1A2, 2C8, and 2D6, the potential is regarded low (with estimated IC_50_ values > 10 μM).

**Figure 8.**
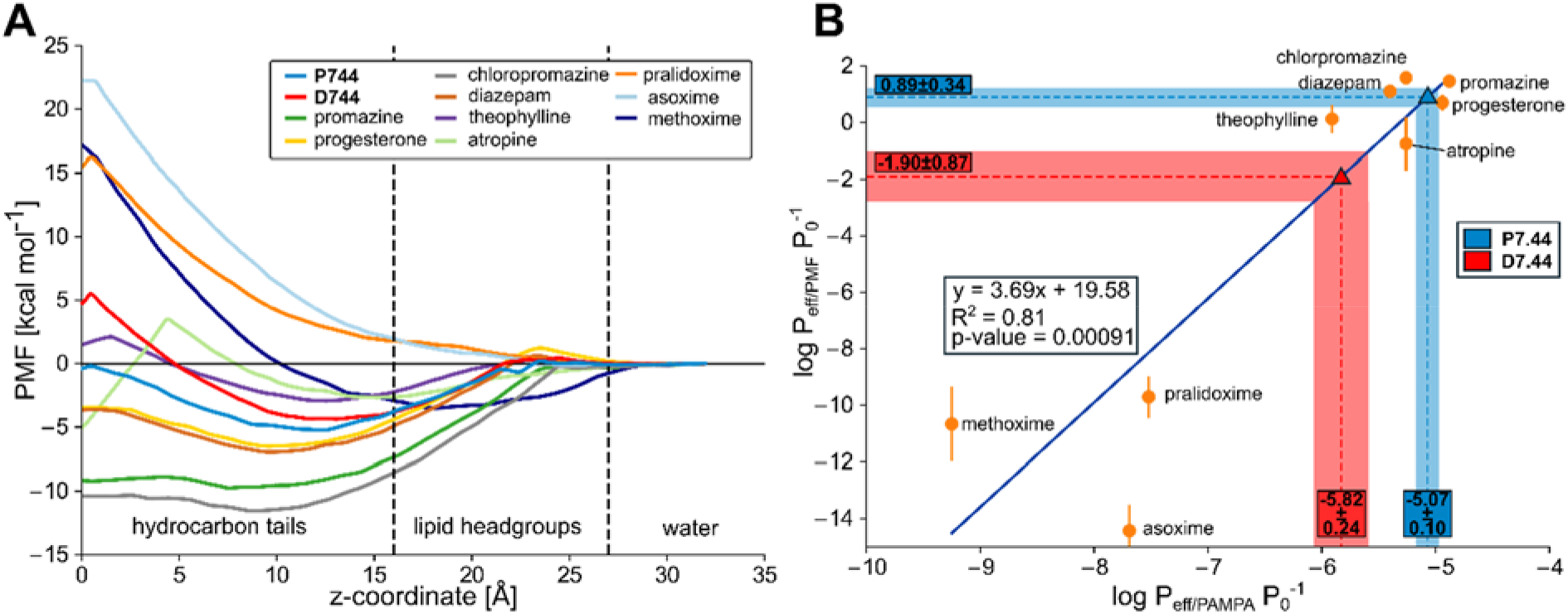
**A)** Potential of mean force (PMF) of permeation of **7.44** in a protonated (P7.44, blue) and deprotonated (D7.44, red) form, and nine calibration molecules. As a reaction coordinate, the distance to the center of a DOPC membrane bilayer was used, setting the profile in the water phase to zero. Molecules with a higher permeability (like progesterone or promazine) display favorable free energy in the membrane center, while those that are impermeable (like the oximes) display a high energy barrier in the membrane center. **B)** Calibration curve between the calculated permeabilities obtained from MD simulations and the experimental values obtained from PAMPA experiments from [58]. In blue (P7.44) and red (D7.44) are the results obtained from the simulations (y-axis, 0.89 ± 0.34 and −1.90 ± 0.87 for P7.44 and D7.44, respectively), and the corresponding predictions obtained from the calibration curve (−5.07 ± 0.10 and −5.82 ± 0.24, respectively), indicating that **7.44** is highly permeable in its protonated state and low to medium permeable in its deprotonated state. *P*_eff_ is the effective permeability, and P_0_ is the unit factor corresponding to 1 cm sec^−1^.

To conclude, **7.44** has favorable physicochemical and ADME properties with high aqueous solubility, high stability in buffer and plasma, and a low hepatic intrinsic clearance *in vitro.* There is a slight potential for CYP-related drug-drug interaction that has to be further investigated.

## Discussion

In this study, we characterized the interaction between the NHR2 domain of RUNX1/ETO, the product of t(8;21) chromosomal translocation that has an oncogenic function in t(8;21) acute myeloid leukemia [6, 10], and the first-in-class molecule to interfere with RUNX1/ETO tetramerization, **7.44**, for targeting acute myeloid leukemia [11, 12]. We established biophysical assays based on DSF and MST to show that **7.44** interferes with NHR2 tetramer stability and increases the dimer population of NHR2, indicating that **7.44** fosters a tetramer-to-dimer transition of NHR2. We also determined the affinity of **7.44** for binding to NHR2; the dissociation constant of **7.44** from the NHR2 dimer, *K*_lig_, is 3.95 ± 1.28 μM. Furthermore, by NMR spectroscopy, we show that **7.44** binds with both heteroaromatic moieties to NHR2 and interacts with or leads to conformational changes in the N-termini of the NHR2 tetramer. Combined with MD simulations of free ligand diffusion, we suggest that at least one of the 1,3-benzodioxoles of **7.44** forms interactions with one of two buried tryptophan residues in the tetramer complex that had been identified as hot spots before. Finally, we demonstrate that the pKa of **7.44** is ~3.9 and that the compound has favorable physicochemical, pharmacokinetic, and toxicological properties.

That **7.44** interacts with the NHR2 domain and interferes with NHR2 tetramer stability was shown in five independent biophysical experiments. First, SEC revealed that the amount of dimeric NHR2 increases with increasing **7.44** concentration, most probably due to a shift in the equilibrium from the tetramer to the dimer. Second, DSF revealed typical curves [16, 61, 62] (**Figure 2A**) applying protein concentrations in commonly used ranges [49, 61, 63, 64]. With increasing **7.44** concentration, the results show decreased fluorescence intensities at the previously determined melting point of the NHR2 tetramer at ~85°C [10] and generally increasing fluorescence intensities at the previously determined melting point of the NHR2 dimer at ~55°C [10]. Third, examining the thermal stability of NHR2 in the absence and presence of **7.44** by CD spectroscopy revealed a cooperative biphasic melting in both cases, with the phase transition at the temperature of dimer melting becoming more pronounced in the presence of **7.44**. These findings demonstrate that **7.44** alters the proportions of the two NHR2 species in a dose-dependent manner towards the dimer and, thus, provide biophysical evidence for the destabilization of NHR2 tetramerization by **7.44**.

Fourth, we quantified the dissociation constant of the NHR2 tetramer into NHR2 dimers (*K*_tet_) and the dissociation constant of **7.44** with respect to the NHR2 dimer (*K*_lig_) using MST, exploiting that the measured molecular movements along a temperature gradient depend on the size, charge, and solvation shell of the molecules [65]. With increasing concentrations, binding of nonfluorescent NHR2 to dye-labeled one resulted in increased fluorescence intensity, whereas binding of **7.44** decreases the fluorescence intensity. From the data, *K*_tet_ = 11.3 ± 1.81 μM was obtained. **7.44** titration data analyzed by three independent modes yielded an *EC*_50_ of 2.5 ± 0.8 μM. As all three analysis modes resulted in very similar *EC*_50_ values, a slow dissociation kinetic of the complex is suggested [19]. According to the obtained Hill coefficients, the binding of **7.44** to NHR2 is not or weakly cooperative, in line with our previous results [11]. In the extreme case of strict binding cooperativity, the Hill coefficient reflects the number of ligand binding sites on a protein [66]; *n* = 1.0-1.5 then indicates a stoichiometry of about one **7.44** molecule per NHR2 monomer. From these results, *K*_lig_ = 3.95 ± 1.28 μM was computed, assuming that **7.44** binds in a 2:1 stoichiometry to the NHR2 dimer and that *K*_lig_ of the first and second ligand-binding event are identical. The assumptions are further supported in that **7.44** shifts the equilibrium from the tetramer to the dimer, the NHR2 dimer is *C*2-symmetric, and the binding sites of the two ligands do not overlap (see below). The *K*_lig_ of **7.44** is comparable to *K*_D_ or *K*_i_ of marketed PPIM or compounds subjected to clinical trials [67], such as pentamidine [68], gossypol [69], or sulindac [70].

Fifth, we used STD-NMR to map the **7.44** epitope binding to NHR2. Multiple protons across **7.44** showed a signal in the STD-NMR spectrum, clearly indicating that **7.44** binds to NHR2, but that not all protons receive the same amount of saturation, indicating that **7.44** interacts with both heteroaromatic moieties with NHR2. These results agree well with the structural model of the **7.44** epitope proposed by us [11] and suggest that modifications at the carboxy group, e.g., a bioisosteric replacement [71] or fluorescent tagging to study in-cell properties [72], should not grossly impact the affinity.

Using multidimensional solution NMR spectroscopy, we further characterized the interaction between **7.44** and NHR2. Observing a good solution NMR spectrum of a coiled-coil domain like NHR2 is highly challenging [54] as the size (34 kDa), rod-like shape (~100 Å length), poor dispersion of the α-helical protein, and large rotational diffusion anisotropy cause poor signal-to-noise ratios in the multiple-resonance spectra [48, 54]. This issue was circumvented by using TROSY-based NMR experiments measured at high temperature on uniformly ^2^H, ^13^C, ^15^N-labeled NHR2 protein. The ^1^H-^15^N TROSY-HSQC spectrum shows one set of peaks for each amino acid of the NHR2 domain, as expected for the highly symmetric homotetramer as shown by X-ray crystallography [48] and the high prevalence of the NHR2 tetramer according to our AUC, SEC, and DLS results. We succeeded in assigning 32% of the backbone amide signals, which are located in the termini of NHR2. Unassigned residues are located in the middle of the coiled-coil region, which is the most rigid NHR2 part.

The interaction of NHR2 with **7.44** was measured via a ^1^H-^15^N TROSY-HSQC spectrum. Pronounced CSP backbone amide resonances (**Figure 5**) indicated binding of **7.44** to NHR2 and/or conformational changes induced by the ligand. CSP were found in the N-terminus (Q485 to L493) and for some unassigned amino acids of the middle region. No CSP were found for the assigned C-terminal residues (Y544 to K552). The C-termini do not form helical interfaces in the tetramer according to the X-ray structure [48]. Of the most affected residues, R492 forms a salt bridge with D533 and a solvent-exposed salt bridge with E536, which interacts with the indole NH of W540 via a charge-assisted hydrogen bond. Notably, residues D533, E536, and W540 were the template motif with which **7.44** was identified by virtual screening and which **7.44** is thought to mimic as a tetramerization inhibitor [11]. The CSP results support this model. Furthermore, CSP of tryptophan side chains indicate the involvement of at least one, if not two, tryptophans in the **7.44** effect.

Next, we performed unbiased MD simulations of free **7.44** diffusion around the NHR2 tetramer to obtain structural information on the binding of the ligand in the context of the oncogenic tetramer, mimicking the situation in the NMR experiments. Such MD simulations have been used previously successfully to identify binding modes of small-molecules at proteins by us [43, 73] and others [74, 75] and complement the NHR2 epitope mapping by NMR by providing information at the atomistic level. Throughout the simulations, the tetramer remained largely structurally invariant (**Figure S13**). Of the three epitopes where **7.44** preferably forms contacts with the NHR2 tetramer, epitopes I and II represent poses in which **7.44** intercalates between the helices of the NHR2 tetramer, once between helices of one dimer and once between helices of two dimers. In either case, **7.44** interacts with at least one of its 1,3-benzodioxoles with tryptophan residues identified as hot spots [11], which may explain the observed STD-NMR results and tryptophan side chain CSP. Although no dissociation of the NHR2 tetramer was observed during our simulation times, in line with the determined *K*_tet_, particularly the intercalation between two helices of two dimers, targeting W540, might represent an early step in the separation of the dimers. Note that this is concordant with our previously proposed binding model for **7.44** at the NHR2 dimer [11, 12]. Still, we could not obtain an (experimentally validated) binding mode of **7.44** at the NHR2 dimer - probably because, on the one hand, the NHR2 dimer population in the NMR experiments was too low ([NHR2] = 4 × *K*_tet_), which shifted the equilibrium to the tetramer side. On the other hand, detecting the precise NHR2 binding mode was hampered because the NHR2 dimer shows marked structural changes at the termini in MD simulations, which makes obtaining converged results for **7.44** binding cumbersome.

Finally, we assessed six physicochemical and *in vitro* pharmacokinetic properties and the pKa of **7.44** experimentally and further pharmacokinetic and toxicological properties computationally, which are key properties in early drug discovery [76]. In all cases, **7.44** shows favorable properties, suggesting that **7.44** is suitably orally bioavailable. Together with its low molecular weight, high ligand efficiency, and non-complex chemical structure [11], this distinguishes **7.44** from other recent first-in-class peptidomimetic protein interaction inhibitors [43].

In summary, we biophysically characterized the interaction between the NHR2 domain of RUNX1/ETO and **7.44** and assessed **7.44**’s physicochemical, *in vitro* pharmacokinetic, and toxicological properties. The results, together with previous biochemical, cellular, and *in vivo* [12] assessments, reveal **7.44** as a lead for further optimization of structurally related compounds with increased binding affinity and enhanced anti-leukemic effects to inhibit RUNX1/ETO oncogenic function in t(8;21) AML.

## Supporting information

Supplemental notes for the cloning, expression, and purification of 6His-NHR2, determination of *K*_tet_ and *K*_lig_ from MST experiments, and permeability estimation from molecular dynamics simulations. Supplemental **Tables S1-S11**. Supplemental **Figures S1 – S13**.

## Acknowledgments

This work was supported by a grant by the state of North-Rhine Westphalia and the European Fonds for Regional Development EFRE.NRW 2014-2020 to HG. We are grateful for computational support and infrastructure provided by the “Zentrum für Informations- und Medientechnologie” (ZIM) at the Heinrich Heine University Düsseldorf and the computing time provided by the John von Neumann Institute for Computing (NIC) to HG on the supercomputer JUWELS at Jülich Supercomputing Centre (JSC) (user ID: HKF7, VSK33). Financial support by Deutsche Forschungsgemeinschaft (DFG) through funds (INST 208/704-1 FUGG to HG) to purchase the hybrid computer cluster used in this study, an Emmy-Noether- and Heisenberg fellowship to ME (ET 103/2 and ET 103/5), and GRK 2158/2 (project number 270650915) to HG is gratefully acknowledged.

