## Supplemental Information for "Biophysical and pharmacokinetic characterization of a small-molecule inhibitor of RUNX1/ETO tetramerization with anti-leukemic effects"

### Supplemental Notes

#### List of abbreviations

| | |
|---|---|
| ACN | acetonitrile |
| AMMC | 3-[2-(*N*,*N*-diethyl-*N*-methylamino)-ethyl]-7-methoxy-4-methyl-coumarin |
| BFC | 7-benzyloxy-trifluoromethyl coumarin |
| CEC | 3-cyano-7-ethoxycoumarin |
| CYP | cytochrome P450 |
| Da | daltons |
| DBF | dibenzylfluorescein |
| DMSO | dimethyl sulfoxide |
| EFC | 7-ethoxy-4-trifluoromethyl-coumarin |
| ESI | electro spray ionization |
| F | fluorescein |
| FA | formic acid |
| FF | furafylline |
| G6PDH | glucose-6-phosphate dehydrogenase |
| H-ESI | heated electrospray ionization (ion source type) |
| HEPES | 4-(2-hydroxyethyl) piperazine-1-ethanesulfonic acid |
| HFBA | heptafluorobutyric acid |
| HPLC | high-performance liquid chromatography |
| HRMS | high-resolution mass spectrometry |
| $IC_{50}$ | concentration inhibiting an enzymatic reaction by 50% |
| ISTD | internal standard |
| KTZ | ketoconazole |
| LC | liquid chromatography |
| m/z | mass-to-charge ratio |
| MeOH | methanol |
| MFC | 7-methoxy-4-trifluoromethyl coumarin |
| MS | mass spectrometry |
| MS-SIM | mass spectrometry-single ion monitoring |
| MW | molecular weight |
| N | normality |
| NaOH | sodium hydroxide |

| | |
|---|---|
| PBS | phosphate-buffered saline (without $Ca^{2+}$ and $Mg^{2+}$) |
| PPB | plasma protein binding |
| pH | potential hydrogen |
| QCT | quercetin |
| QND | quinidine |
| Q-TOF | quadrupole time-of-flight |
| RFU | relative fluorescence unit |
| RT | room temperature or retention time |
| RV | recovery |
| S | seconds |
| SFZ | sulfaphenazole |
| S/N | signal-to-noise ratio |
| TCP | tranylcypromine |
| Tris HCl | tris(hydroxymethyl)-amino-methan hydrochlorid |
| v/v | volume per volume |
| v | volts |
| w/v | weight per volume |
| WME | William's medium E |

**Cloning, expression, and purification of 6His-NHR2**

Synthetic DNA corresponding to the coding region of residues 498-548 of RUNX1/ETO was cloned into expression vector pET19b in *E. coli* BL21 (DE3). A cysteine-free version of the NHR2 sequence was used where both cysteines were replaced by serines to comply with the construct used in previous studies [1, 2].

A His-tag was added to the N-terminus to allow for purification of NHR2 via immobilized metal ion affinity chromatography (IMAC). The transformed *E. coli* was transferred into LB-medium (5 g yeast extract, 5 g NaCl, and 10 g tryptone or peptone), containing ampicillin, and incubated at 37 °C until an $OD_{600}$ of ~1 was reached. The expression was induced by adding (per liter LB-medium) 1 ml 1 M isopropyl-1-thio-β-D-galactopyranoside. The expression was continued for ~4 h at 37 °C. The cells were harvested by centrifugation and disrupted via sonification. The target protein was purified using IMAC, and the purity of the protein was inspected using a 15% SDS gel (Figure S1)

**Determination of $K_{tet}$ and $K_{lig}$ from MST experiments**

The $EC_{50}$ value determined by MST in this study as well as the $EC_{50}$ values determined by $BS^3$ cross-linking assay and ELISA previously [3] are composites of the dissociation constant of the NHR2 tetramer ($K_{\text{tet}}$) and the dissociation constant of **7.44** to the NHR2 dimer ($K_{\text{lig}}$). As the NHR2 tetramer (T) is in equilibrium with its dimeric form (D), the ligand (L) can only bind to the NHR2 dimer (D) after dissociation of T. Given the symmetry of an NHR2 dimer, a second ligand can bind to the DL complex, resulting in a DLL complex (see eq. 4 in the main text). The respective equilibrium constants are defined as:

$$K_{\text{tet}} = [D]^2 / [T] \tag{S1a}$$

and

$$K_{\text{lig}} = [D] \cdot [L] / [DL]. \tag{S1b}$$

$K_{\text{lig}}$ could not be determined directly. Instead, $K_{\text{lig}}$ was determined as a function of $EC_{50}$ and $K_{\text{tet}}$ according to the following equations [4, 5]. The following derivation was taken from ref. [6] and has been applied in a related context in ref. [4].

In the absence of L, the concentration of unbound NHR2 dimer D is $[D]_0$, and the concentration of the NHR2 tetramer T is $[T]_0$. The total concentration of NHR2 dimer $[D]_{\text{tot}}$ is (eq. S2):

$$[\mathrm{D}]_{\mathrm{tot}} = 2\,[\mathrm{T}]_0 + [\mathrm{D}]_0 \tag{S2}$$

Substituting eq. S2 into eq. S1a and solving the obtained equation with respect to $[D]_0$ leads to eq. S3:

$$[\mathrm{D}]_0 = -\frac{\mathrm{K_{tet}}}{4} + \sqrt{\left(\frac{\mathrm{K_{tet}}}{4}\right)^2 + \frac{\mathrm{K_{tet}}[\mathrm{D}]_{\mathrm{tot}}}{2}} \tag{S3}$$

Substituting eq. S3 into eq. S2 and rearranging leads to eq. S4, allowing the calculation of the maximal NHR2 tetramer concentration $[T]_0$ as a function of $K_{tet}$ and $[D]_{tot}$.

$$[\mathrm{T}]_0 = \frac{[\mathrm{D}]_{\mathrm{tot}} + \frac{\mathrm{K_{tet}}}{4} - \sqrt{\left(\frac{\mathrm{K_{tet}}}{4}\right)^2 + \frac{\mathrm{K_{tet}}[\mathrm{D}]_{\mathrm{tot}}}{2}}}{2} \tag{S4}$$

In the presence of a total ligand concentration of $[L]_{tot}$ = $EC_{50}$, the total NHR2 dimer concentration $[D]_{tot}$ is defined by eq. S5:

$$[\mathrm{D}]_{\mathrm{tot}} = 2[\mathrm{T}]_{50} + [\mathrm{D}]_{50} + [\mathrm{DL}]_{50} + [\mathrm{DLL}]_{50} \tag{S5}$$

in which $[T]_{50}$, $[D]_{50}$, $[DL]_{50}$, and $[DLL]_{50}$ are the concentrations of the respective molecular species at $[L]_{tot} = EC_{50}$.

MST experiments in the presence of **7.44** were performed at an NHR2 concentration where >99% of NHR2 is in a dimeric arrangement. Neglecting the tetramer and assuming that the majority of the MST signal change comes from the D to DL equilibrium, $[D]_{50}$ is half of the maximal dimer concentration $[D]_0$ at [L] = 0 (eq. S6).

$$[\mathrm{D}]_{50} = \frac{[\mathrm{D}]_0}{2} \tag{S6}$$

The concentration of the unbound NHR2 tetramer $[T]_{50}$ is obtained by substituting eq. S6 into eq. S1a (eq. S7):

$$[\mathrm{T}]_{50} = \frac{[D]_{50}^2}{K_{tet}} = \frac{[D]_0^2}{4K_{tet}} \tag{S7}$$

Based on the definition of the dissociation constant of the **7.44**-bound complexes $K_{lig}$, the concentrations of the ligand-bound complexes are given by eq. S8 and eq. S9

$$[DL]_{50}=\frac{[D]_{50}[L]_{50}}{K_{lig}} \tag{S8}$$

$$[DLL]_{50}=\frac{[D]_{50}[L]_{50}^2}{K_{lig}^2} \tag{S9}$$

if one assumes that the dissociation constants for both ligand binding events are equal, in agreement with the remote ligand binding sites and our model (eq. 4) that depends on the pre-dissociation of T. $[L]_{50}$ is the concentration of the unbound PPIM at $EC_{50}$.

Substituting eq. S8 and eq. S9 into eq. S5 and solving the resulting quadratic equation with respect to $K_{lig}$ leads to eq. S10. The negative solution of eq. S10 has no physical meaning because of $K_{lig} \geq 0$ and was omitted.

$$K_{lig}=\frac{-[D]_{50}[L]_{50}}{2(2[T]_{50}+[D]_{50}-[D]_{tot})}+\frac{1}{2}\sqrt{\left(\frac{[D]_{50}[L]_{50}}{(2[T]_{50}+[D]_{50}-[D]_{tot})}\right)^2-\frac{4[D]_{50}[L]_{50}^2}{(2[T]_{50}+[D]_{50}-[D]_{tot})}} \tag{S10}$$

According to eq. S6 and eq. S7, the terms $[D]_{50}$ and $[T]_{50}$ in eq. S10 are constants for given $[D]_0$ and $K_{tet}$. Thus, eq. S10 is a function of $[L]_{50}$, which remains to be defined.

For $[L]_{tot} = EC_{50}$, the total ligand concentration $[L]_{tot}$ is defined by eq. S11.

$$[L]_{tot}=EC_{50}=[L]_{50}+[DL]_{50}+2[DLL]_{50} \tag{S11}$$

Substituting eq. S8 and eq. S9 into eq. S11 and solving the resulting quadratic equation with respect to $[L]_{50}$ leads to eq. S12. The negative solution of eq. S12 has no physical meaning because of $[L]_{50} \geq 0$ and was omitted.

$$[L]_{50}=-\frac{\left(\frac{[D]_{50}}{K_{lig}}+1\right)}{4\frac{[D]_{50}}{K_{lig}^2}}+\sqrt{\left(\frac{\left(\frac{[D]_{50}}{K_{lig}}+1\right)}{4\frac{[D]_{50}}{K_{lig}^2}}\right)^2+\frac{EC_{50}}{2\frac{[D]_{50}}{K_{lig}^2}}} \tag{S12}$$

According to eq. S7, $[D]_{50}$ in eq. S12 is constant for given $[D]_0$. Finally, solving the system of eq. S10 and eq. S12 allows calculating $K_{lig}$ for given $[D]_{tot}$, $K_{tet}$, and $EC_{50}$. For this purpose, $[D]_{tot}$ is the total concentration of NHR2 dimer in the assay.

### *In vitro* ADME investigations of compound 7.44

The materials and methods of the six assays, aqueous solubility, plasma protein binding (PPB), plasma stability, hepatocyte clearance measurements, chemical stability, and cytochrome P450 (CYP) inhibition, are described in detail below [7-12].

#### Materials

In the assays described herein, the materials were used as shown in **Table S1**.

**Table S1**. Details of materials.

| Material | Catalog ID | Supplier | Lot |
|---|---|---|---|
| **Internal standards** | | | |
| Diazepam | D0899 | Sigma-Aldrich | 105F0451 |
| Diclofenac | D6899 | Sigma-Aldrich | BCBN3367V |
| Griseofulvin | PHR1534 | Sigma-Aldrich | LRAB4111 |
| **Solubility** | | | |
| PBS (pH 7.4) | P04-36500 | PAN Biotech | 1516119 |
| **PPB** | | | |
| Mouse plasma (CD-1, male, pooled, Li-heparin) | MSEPLLIHP-M | Biotrend | MSE308232 |
| Warfarin | 45706 | Sigma-Aldrich | SZBF197XV |
| **Plasma stability** | | | |
| Mouse plasma (CD-1, male, pooled, Li-heparin) | MSEPLLIHP-M | Biotrend | MSE308232 |
| Propantheline | P8891 | Sigma-Aldrich | 110M1921 |
| **Chemical stability** | | | |
| PBS (pH 7.4) | P04-36500 | PAN Biotech | 1560119 |
| Omeprazole | O104 | Sigma-Aldrich | BCBQ8476V |
| **Hepatocyte stability** | | | |
| 7-Ethoxycoumarin | E1379 | Sigma-Aldrich | MKBN6201V |
| 1 M HEPES solution | P05-01100 | PAN Biotech | 2190518 |
| In vitro GRO HT medium | Z99019 | IVT Bioreclamation | C11020A |
| L-glutamine | P04-80100 | PAN Biotech | 4140319 |
| Primary mouse hepatocytes[a] | M005052 | IVT Bioreclamation | MHD |
| Williams Medium E | P04-29510 | PAN Biotech | 2861019 |
| **CYP inhibition using HITS™ kits** | | | |
| Furafylline | F124 | Sigma-Aldrich | 59790 |
| Tranylcypromine[b] | P8511 | Sigma-Aldrich | MKBW0574V |
| Quercetin dihydrate | 0020-05-95 | Sigma-Aldrich | HWI00580-2 |
| Sulfaphenazole | S0758 | Sigma-Aldrich | 63530 |
| Quinidine HCl monohydrate | Q0750 | Sigma-Aldrich | SLSS3542V |
| Ketoconazole | K1003 | Sigma-Aldrich | SLCC4160 |

| | | | |
|---|---|---|---|
| G6PDH | G5885 | Sigma-Aldrich | 096K8605 |
| CYP1A2/CEC kit | 459500 | Corning | 5239001 |
| CYP2B6/EFC kit | 459220 | Sigma-Aldrich | 4296001 |
| CYP2C8/DBF kit | 459320 | Corning | 4300001 |
| CYP2C9/MFC kit | 459300 | Corning | 535003 |
| CYP2C19/CEC kit | 459400 | Corning | 5313006 |
| CYP2D6/AMMC kit | 459200 | Corning | 5162021 |
| CYP3A4/BFC kit | 459100 | Corning | 6102001 |

[a] cryopreserved, strain ICR/CD-1, male, pooled

[a] *trans*-2-phenylcyclopropyl-aminehydrochloride

**Buffers / media / solvents**

In the assays described herein, solvents were used as shown in **Table S2**.

**Table S2**. Solvents

| Material | Catalog ID | Supplier |
|---|---|---|
| Acetonitrile (ACN) | 34851-PPB | Honeywell |
| Deionised water ($H_2O$) | NANOpure DIamond Life Science Water purification system | |
| Dimethylsulfoxide (DMSO) | 4720.2-PPB | Carl Roth |
| Ethanol (EtOH) | 32205 | Honeywell |
| Methanol (MeOH) | 34885 | Honeywell |

Further buffers were used in the assays:

**Solubility in aqueous solutions**:

Ready to use PBS, pH 7.4, as described in **Table S1** was used.

**Hepatocyte stability assay**:

InvitroGRO HT Medium: Ready-to-use medium for thawing of cells was applied.

Incubation medium: WME supplemented with 25 mM HEPES and 2 mM L-glutamine

**Description of key quantitative analytical equipment**

LC-MS: Surveyor MS Plus HPLC (Thermo Electron) HPLC system connected to a TSQ Quantum Discovery Max (Thermo Electron) triple quadrupole mass spectrometer equipped with an electrospray (ESI) (Thermo Fisher Scientific, USA); connected to a PC running the standard software Xcalibur 2.0.7. It is used for solubility, PPB, and plasma stability assays.

LC-MS: 1260 Infinity II BioInert Quarternary U-HPLC pump, 1260 Inifinity II BioIntert Multisampler, 1260 Inifinity II BioIntert Column Compartment and 1260 Inifinity II Diode Array Detector (Agilent, Germany) connected to a QTOF X500B (SCIEX, Canada) mass spectrometer, data handling with the standard software SCIEX OS (Version 1.5 and higher). It is used for chemical stability and hepatocytes stability assays.

Fluorimeter: Wallac Victor 1420 Multilabel Counter, Perkin Elmer.

**Methods**

**Preparation of test solutions**

In **Table S3**, the concentrations of the test and reference item stock solutions in their respective solvents are displayed. By default, working solutions of test and reference items were diluted from the stock solutions in an appropriate solvent to obtain working solutions of 100-fold (solubility), 50-fold (PPB, plasma stability, chemical stability), and 200-fold (hepatocyte stability) higher strength than the final incubation concentrations and the respective final organic solvent content (**see next section**).

**Table S3**. Stock and working solutions of test and reference items

| **Item** | **Stock solution** | | **Working solution solvent** |
|---|---|---|---|
| | (mM) | Solvent | |
| **7.44** compound | 20 | DMSO | DMSO (Solubility, Hepatocyte stability) 25% DMSO in $H_2O$ (PPB, Plasma stability, Chemical stability) |
| Propantheline (Plasma stability) | 10 | $H_2O$ | $H_2O$ |
| Omeprazole (Chemical stability) | 10 | DMSO | 25% DMSO in $H_2O$ |
| 7-Ethoxycoumarin (Hepatocyte stability) | 10 | ACN | ACN |

**Test concentration selection**

The test item **7.44** compound was dissolved in DMSO at 20 mM. The cut-off concentration of test item **7.44** in the **solubility assay** was 200 µM with a final solvent content of 1% DMSO. **For PPB assay**, the test item was tested at final concentrations of 10 µM in the presence of 0.5% DMSO. Reference item warfarin was tested at a standardized final concentration of 10 µM in the presence of 1% ACN.

**For plasma stability assay**, working solutions of the test and reference item were diluted from the stock solutions in the appropriate solvent (**Table S3**) to reach working solutions of 50-fold higher strength than the final intended incubation concentrations of the compounds. Test item and propantheline (reference item for low plasma stability) were tested at a final concentration of 10 µM. The final solvent content in the test item incubations was 0.5% DMSO, while in the propantheline incubations, no organic solvent was added.

**For chemical stability assay**, working solutions of the test and reference items were diluted from the stock solutions in the appropriate solvent (**Table S3**) to reach working solutions of 50-fold higher strength than the final intended incubation concentrations of the compounds. Test item and Omeprazole (reference item for low chemical stability) were tested at a final concentration of 10 µM. The final solvent content in the test item incubations was 0.5% DMSO.

**For the hepatic stability assay**, working solutions of the test and reference items were diluted from the stock solutions in the appropriate solvent (**Table S3**) to reach working solutions of a 200-fold higher strength than the intended final concentrations. In a second step, working solutions were further diluted in incubation medium to reach 10-fold concentrated starting solutions. The final test concentrations were 5 µM with a solvent content of 0.5% DMSO or 1% ACN in all test item and reference item incubations, respectively.

**For CYP inhibition assay**, the test item was tested at final concentrations of 1 µM and 10 µM, and reference items were tested at eight appropriate concentrations. Stock solutions of test items were diluted in DMSO to prepare a 200-fold concentrated working solution for CYP2B6, or stock solutions were diluted in NADPH-cofactor-mix (all other CYPs) to reach concentrations 20-fold higher than the requested final test concentration of 10 µM and 1 µM, respectively. The final organic solvent content was 0.5 % DMSO for CYP2B6 or 0.2% DMSO for all other tested CYP isoforms, respectively. Stock solutions of the reference items were diluted 50-fold in the first well of a row for CYP1A2, CYP2C9, CYP2C19, CYP2D6, and CYP3A4, 100-fold for CYP2C8, 200-fold for CYP2B6, followed by a serial 1:3 dilution in the 96-well plates containing cofactor mix to obtain the respective eight test concentrations. The final solvent content in the incubations was 2% ACN for CYP1A2, CYP2C9, CYP2C19, CYP2D6, CYP3A4 and 1% ACN for CYP2C8 and 0.5% ACN for CYP2B6.

**Preparation of calibration standards of test and reference items**

By default, working solution of the test and reference items were prepared for each calibration level. The calibration standards were ranged from 0.02 µM to 300 µM for the solubility assay, from 0.001 µM to 20 µM for the PPB assay, from 0.05 µM to 20 µM for the plasma stability and chemical stability assays, and from 0.005 µM to 10 µM for the hepatocytes stability assay by appropriate dilution of the corresponding stock solution with the corresponding solvent by serial dilution.

**Solubility**

Calibration standards for the solubility assay were prepared by spiking 198 µL of the respective buffer (PBS buffer (pH 7.4)) with 2 µL of the corresponding working solution. The final standard solutions contained 1% DMSO.

**PPB**

For the PPB assay calibration standards were prepared by spiking 73.5 µL mouse plasma with 1.5 µL of the corresponding working solution. To avoid unspecific test item metabolism in calibration standards, protein was inactivated by addition of 150 µL ACN containing the internal standards (Diazepam, 1 µM, Griseofulvin, 1 µM, and Diclofenac, 10 µM) prior to addition of working solutions. The final standard solutions contained 1% DMSO (test item) or 1% ACN (reference item).

**Plasma stability and Chemical stability**

Calibration standards for plasma stability and chemical stability assays were prepared by spiking 147 µL mouse plasma or PBS buffer, respectively with 3 µL of the corresponding working solution. Plasma was inactivated by addition of 300 µL ACN containing the internal standards (Diazepam, 1 µM, Griseofulvin, 1 µM, and Diclofenac, 10 µM) before addition of working solutions to avoid unspecific metabolism in calibration standards. The final standard solutions contained 0.5% DMSO.

**Hepatocyte stability assay**

Calibration standards were prepared by mixing the appropriate volumes of incubation medium (196 µL) and the corresponding working solution (4 µL). The final standard solutions contained 2% DMSO or 1% ACN for test item and reference item, respectively.

Calibration solutions were processed for precipitation (ACN containing the internal standards) and quantitative bioanalysis as described in **Sample preparation section**.

**Assay procedures**

**7.44** compound was prepared as 20 mM stock solution in DMSO and further diluted in 25% DMSO/$H_20$ or assay matrix if applicable.

**Kinetic solubility in PBS**

2.5 µl of the compound **7.44** stock solution was mixed with 247.5 µl of PBS (pH 7.4), resulting in a cut-off concentration of 200 µM and a final solvent content of 1% DMSO. The test solutions (triplicates) were put on a filter plate and were shaken protected from light at 300 rpm at room temperature for 1.5 hours, followed by centrifugation at 500 x g for 3 minutes. 200 µl of the filtrate were mixed with 100 µl ACN containing the ISTD as described in **Sample preparation section**.

**PPB**

Plasma protein binding was performed according to the modified ultrafiltration procedure described by Taylor and Harker[12]. Test and reference item stock solutions were diluted in 25% DMSO/$H_2O$ or ACN, respectively (see **Preparation of test solutions section**) to give a working solution of 50-fold higher strength than the intended final test concentration. The plasma was prewarmed for 30 min at 37°C. The incubation solutions were prepared by adding 1.5 µL of the 50-fold concentrated working solution to 73.5 µL of plasma and vortexed for 2 min, resulting in a final incubation concentration of 10 µM for the test items (0.5% DMSO final solvent concentration) and of 10 µM for the reference item warfarin in the presence of 1% ACN. The assay was performed with plasma from CD-1 mice (**Materials section**). Plasma samples (with test or reference item) as well as the control plasma samples (without test or reference item) were incubated at 37°C for 60 min in the dark.

After incubation, the plasma samples (with test or reference item) were added to sample reservoirs of Microcon® centrifugal filter units (Ultracel-YM30, MWCO 30000 Da; Millipore, USA) For each unit loaded with plasma sample, a partner ultrafiltration unit was loaded with control plasma. All ultrafiltration units were centrifuged (6,500 x g for 12 minutes, room temperature). The sample reservoirs containing plasma retentate were then inverted and placed on the filtrate collection tubes of the partner ultrafiltration unit. The ultrafiltration units were centrifuged a second time (700 x g for 20 seconds), such that the retentate was mixed with the filtrate of the partner sample. As a result, two reconstituted plasma samples were produced, one representing drug in the filtrate and one representing drug in the retentate. The reconstituted samples were processed for sample preparation as described below.

In parallel, stability control plasma samples (with test or reference item) were prepared and incubated for 0 or 60 min, respectively. After incubation, stability control plasma samples were processed for sample preparation as described below to determine the post-incubation test item concentration.

All experimental incubations containing the test items were run in duplicate (n = 2). Warfarin was used as high-binding positive control (n = 3).

**Plasma Stability and Chemical Stability**

The test item stock solution was diluted in 25% DMSO/$H_2O$ to prepare a working solution of 50-fold higher strength than the intended incubation concentration and kept protected from light. The incubation solutions were prepared by adding 3 µL of this 50-fold concentrated working solution to 147 µL of pre-incubated assay matrix (i.e. PBS (pH 7.4) or plasma pre-warmed to 37°C for chemical stability and plasma stability assays, respectively) resulting in an incubation concentration of 10 µM. Sampling time points were (0, 30, 60, 120, and 240 minutes) for plasma stability assay and (0 and 240 minutes) for chemical stability assay. Per time point, two replicates were conducted. The final solvent concentration was 0.5% DMSO for the test items. The plasma stability and chemical stability assays were performed using plasma from CD-1 mice and PBS buffer (pH 7.4), respectively.

Propantheline, a hydroxyethyl-diisopropylmethyl-ammonium xanthene-9-carboxylate, is cleared from plasma rapidly by the release of the hydroxyl-ethyl-diisopropylmethylammonium moiety catalyzed by esterases and was included as positive control (n = 2) in the experiment. These results were used as marker for comparison of test item reactivity in plasma.

For the positive control propantheline, samples were removed after 0 and 120 minutes of incubation with mouse plasma. Samples were stopped by addition of ACN containing ISTDs and processed.

Omeprazole is chemically unstable and easily degradable in buffer, so it was included as positive control (n = 2) in the experiment. Samples were removed after 0 and 240 minutes of incubation of omeprazole samples with PBS buffer. The reaction was stopped by addition of ACN containing ISTDs and processed as described in **Sample preparation section**.

**Hepatocyte stability**

Working solutions of the test and reference items were diluted from the stock solutions in the appropriate solvent (**Table S3**) to reach working solutions of a 200-fold higher strength than the intended final concentrations. In a second step, working solutions were further diluted in incubation medium to reach 10-fold concentrated starting solutions. The final test concentrations were 5 μM with a solvent content of 0.5% DMSO or 1% ACN in all test item and reference item incubations, respectively. **7.44** compound was dissolved 20 mM in DMSO and 25% DMSO/$H_2O$. Primary hepatocytes from CD-1 mice (pooled, male) were thawed according to the instructions of the manufacturer. The incubation samples were composed of 0.2 x $10^6$ cells/well in 225 μL incubation medium and 25 μL test item solution (50 μM in incubation medium), resulting in a final start concentration of 5 μM. Samples were taken from the suspension cultures after 0, 15, 30, 60, and 120 minutes of incubation and processed for LC-MS analysis as described in **Sample preparation section**.

Positive control incubations were performed using 7-ethoxycoumarin as substrate. Metabolic turnover rates were measured at 0 and 120 minutes of incubation. Aliquotes were taken from the incubations for sample preparation and analysis. Hepatocyte enzyme activity was assessed in terms of 7-ethoxycoumarin turnover, i.e., loss of 7-ethoxycoumarin.

Negative controls were performed to observe non-metabolic degradation processes; i.e. test item concentrations remaining stable over the investigated time suggests that a decrease of the parent compound is mainly due to metabolism. Negative control incubations were performed in line with all experiments using incubation medium with test and reference item in the absence of hepatocytes. Samples were taken from the incubations at 0 and 120 minutes and processed as described below (in **Sample preparation section**).

**CYP inhibition using HTS™ kits**

Assays were performed in black 96-well plates according to the manufacturer's instructions (BD Gentest/Corning, P450 High Throughput Inhibitor Screening Kits). The final concentrations of the reagents required for the individual CYP isozymes are given in **Table S4**. Shortcuts given in the table are shown below.

For determination of reference item $IC_{50}$ values, twelve wells in one row including solvent control (100% enzyme activity, no inhibition) and negative controls (NC, blank values) were used for inhibition curve for each reference item. All reactions were performed in duplicates. The source of NADPH in these enzyme assays was a NADPH-regenerating system: glucose-6-phosphate-dehydrogenase converts $NADP^+$ to NADPH in the presence of glucose-6-phosphate.

Cofactor mix, containing the $NADP^+$-regenerating system, was prepared according to the manual and was filled in the plate to well 1 for determination of $IC_{50}$ values of reference items or to each test item well. Cofactor/solvent mix was prepared according to the manual and was filled in the plate from well 2 to well 12 of the reference items, and in solvent control as well as in NC wells for test items. The working solutions of the test items were added to each test well.

For reference items, working solutions were diluted 50-fold in the first well of a row for CYP1A2, CYP2C9, CYP2C19, CYP2D6, and CYP3A4, 100-fold for CYP2C8 or 200-fold for CYP2B6, followed by a serial 1:3 dilution from well 1 to 8. Wells 9 and 10 did not contain inhibitor (solvent control, 100% enzyme activity, no inhibition). For the test items, the cofactor mix-containing working solutions were diluted 1:10 (v/v) in the respective wells. All wells contained final organic solvent concentrations as given in **Test concentration selection** section.

**Table S4**. Final concentrations in P450 inhibition screening according to the manufacturer[a]

| Isoenzyme | CYP1A2 | CYP2B6 | CYP2C8 | CYP2C9 |
|---|---|---|---|---|
| Substrate | CEC, | EFC | DBF | MFC |
| Substrate concentration | 5 µM | 2.5 µM | 1 µM | 75 µM |
| Metabolite formed | CHC | HFC | F | HFC |
| NADP+ | 1.3 mM | 1.3 mM | 1.3 mM | 1.3 mM |
| Glucose-6-phosphate | 3.3 mM | 3.3 mM | 3.3 mM | 3.3 mM |
| $MgCl_2$ x $6H_2O$ | 3.3 mM | 3.3 mM | 3.3 mM | 3.3 mM |
| Glucose-6-phosphate-dehydrogenase | 0.4 U/mL | 0.4 U/ml | 0.4 U/ml | 0.4 U/ml |
| Phosphate buffer, pH 7.4 | 100 mM | 100 mM | 50 mM | 25 mM |
| Enzyme | 2.5 pmol/ml | 5 pmol/ml | 20 pmol/ml | 5 pmol/ml |
| Reference inhibitor, highest concentration | FF 100 µM | TCP 5 µM | QCT 20 µM | SFZ 10 µM |

| Isoenzyme | CYP2C19 | CYP2D6 | CYP3A4 |
|---|---|---|---|
| Substrate | CEC | AMMC | BFC |
| Substrate concentration | 25 µM | 1.5 µM | 50 µM |
| Metabolite formed | CHC | AHMC | HFC |
| NADP+ | 1.3 mM | 8.2 µM | 1.3 mM |
| Glucose-6-phosphate | 3.3 mM | 0.41 mM | 3.3 mM |
| MgCl2 x 6H2O | 3.3 mM | 0.41 mM | 3.3 mM |
| Glucose-6-phosphate-dehydrogenase | 0.4 U/ml | 0.4 U/ml | 0.4 U/ml |
| Phosphate buffer, pH 7.4 | 50 mM | 100 mM | 200 mM |
| Enzyme | 2.5 pmol/ml | 7.5 pmol/ml | 5 pmol/ml |
| Reference inhibitor, highest concentration | TCP 100 µ M | QND 0.5 µM | KTZ 5 µM |

[a]AHMC = 3-[2-(*N*,*N*-diethylamino)ethyl]-7-hydroxy-4-methylcoumarin AMMC = 3-[2-(*N*,*N*-diethyl-*N*-methylamino)ethyl]-7-methoxy-4-methylcoumarin, BFC = 7-benzyloxy-trifluoromethyl coumarin, C = coumarin, CEC = 3-cyano-7-ethoxycoumarin CHC = 3-cyano-7-hydroxycoumarin, DBF = dibenzylfuorescein, DDTC = diethyldithiocarbamic acid, EFC = 7-ethoxy-4-trifluoromethylcoumarin, F = fluorescein, FF = furafyllin, HC= 7-hydroxycoumarin, HFC = 7-hydroxytrifluoromethyl coumarin, KTZ = ketoconazole, MFC = 7-methoxy-4-trifluoromethyl coumarin, QCT = quercetin, QND = quinidine, SFZ = sulfaphenazole, TCP = tranylcypromine.

After 10 minutes of pre-incubation at 37°C, the reactions were started by addition of pre-warmed enzyme/substrate mix. Incubations with a final volume of 200 µl per well were performed for 15 minutes (CYP1A2), 30 minutes (CYP2B6, CYP2C19, CYP2D6, CYP3A4 – BFC substrate), 40 minutes (CYP2C8) or 45 minutes (CYP2C9) at 37°C. The reaction was stopped as described below (in **Sample preparation**).

Test item was tested for auto-fluorescence, which might interfere with the measurement of the fluorescent metabolite. For this purpose, a solution, replacing cofactor-mix and enzyme/substrate-mix was prepared by mixing appropriate volumes of assay buffer, control protein, and water (auto-fluorescence control mix, AFC). An aliquot of AFC mix was removed and spiked with the respective solvent at the concentration suitable for the individual isoenzyme. 90 µl of the AFC/solvent mix was filled in the plate for all test item wells.

10 µl of the test item working solutions, concentrated as described in **Test concentration selection** section, were added to each AFC well in duplicate for each test concentration (1 µM and 10 µM). The plate was incubated for 10 min at 37°C, imitating the pre-incubation step of test item in cofactor mix. At the end of the pre-incubation, 100 µl of AFC mix was added to each well. Incubations with a final volume of 200 µl per well were performed as described above.

The fluorescence was measured using a Wallac Victor fluorescence plate reader, and the wavelengths for excitation and emission were shown in **Table S8**.

**Sample preparation**

As ISTDs for LC-MS analysis, compounds were chosen from the Pharmacelsus pool known to be suitable for ACN precipitation. Injection volumes of all measurements are listed in **Table S5**.

**Kinetic solubility in PBS**

The sample solution preparation was performed by mixing the 0.5-fold volume of ACN containing the internal standards (1 µM Diazepam, 1 µM Griseofulvin, and 10 µM Diclofenac) with the sample (i.e., filtrate) or calibration standard solution. After vigorously shaking (10 seconds), the samples were centrifuged (2200 x g) for 5 minutes at room temperature. Aliquots (70 µL) of the particle-free supernatants were transferred to 200 µl sample vials and subsequently subjected to LC-MS/MS.

**PPB**

The sample and standard preparation for the PPB assay were performed by mixing 150 µl of ACN containing the internal standards (1 µM Diazepam, 1 µM Griseofulvin, and 10 µM Diclofenac) with 75 µl sample or calibration standard solutions. After vigorously shaking (10 seconds), the samples were centrifuged (6,800 x g) for 5 minutes at room temperature. Particle-free supernatants were diluted with an equal volume of $H_2O$ to reduce the organic solvent content of the samples to 33% and subsequently subjected to LC-MS analysis.

**Plasma Stability and Chemical Stability**

Isolation of the compounds was performed by addition of 300 µl ACN containing the internal standards (1 µM Diazepam, 1 µM Griseofulvin, and 10 µM Diclofenac) to 150 µl samples and calibration standard solutions. After shaking (10 seconds) and sonification (10 seconds), the samples were centrifuged (2200 x g) at room temperature for 10 min. Aliquots (80 µl) of the particle-free supernatants were diluted with an equal volume of PBS buffer to reduce the organic solvent content to 33%. The resulting samples were transferred to a 96-well plate and subsequently subjected to LC-MS/MS.

**Hepatocyte stability**

Metabolic stability samples from hepatocyte incubations were stopped by addition of 200 µl ACN containing the internal standards (1 µM Diazepam, 1 µM Griseofulvin, and 10 µM Diclofenac) to 200 µL sample or calibration standards. Samples were shaken vigorously (10 seconds) then centrifuged (4800 x g, room temperature, 5 minutes). Aliquots of the particle-free supernatants (100 µL) were diluted with an equal volume of $H_2O$ to reduce the organic solvent content of the samples to 25%. The resulting sample was transferred to autosampler vials and subsequently subjected to LC-MS analysis.

**CYP inhibition using HTS$^{TM}$ kits**

For CYP2C8, the reaction was stopped by addition of 75 µl 2N NaOH. Before analysis, samples were further incubated at 37°C for 2 h to increase the signal-to-noise ratio.

The reactions of all other CYP isoforms were stopped by the addition of 75 µl/well stop solution containing 60% ACN and 40% 0.1 M Tris, pH 9.0. To wells 11 and 12, the stop solution was added prior to the addition of enzyme/substrate mix to serve as blanks for background fluorescence.

**Table S5**. Injection volumes.

| **Analyte** | **Injection volume (µl)** |
|---|---|
| Solubility | |
| **7.44** compound | 20 |
| PPB | |
| **7.44** compound | 10 |
| Plasma stability | |
| **7.44** compound | 9 |
| Propantheline | 3 |
| Chemical stability | |
| **7.44** compound | 15 |
| Omeprazole | 15 |
| Hepatocyte stability | |
| **7.44** compound | 15 |

**Key instruments**

**Liquid chromatography – mass spectrometry (LC-MS)**

For quantitative analysis of test and reference items, LC-MS systems as described in **Description of key quantitative analytical equipment** section were used. The pump flow rate was set to 600 µL/min and the analytes were separated on a Kinetex Phenyl-Hexyl analytical column, 2.6 µm, 50 x 2.1 mm (Phenomenex, Germany) with a corresponding pre-column using the gradients as presented in **Table S6**.

For solubility, PPB, and plasma stability measurements applying the triple quadrupole technology, full scan mass spectra were acquired in the positive mode using syringe pump infusion to identify the protonated quasimolecular ions $[M+H]^+$. Auto-tuning was carried out for maximizing ion abundance followed by the identification of characteristic fragment ions using a generic parameter set: ESI ion-transfer-capillary temperature 350°C, capillary voltage 3.8 kV, collision gas 0.8 mbar argon, sheath gas, ion sweep gas, and auxiliary gas pressure were

40, 2, and 10 (arbitrary units). Ions with the highest S/N ratio were used to quantify the item in the selected reaction monitoring mode (SRM) and as qualifier, respectively.

For chemical and hepatocytes stability measurements applying the Q-TOF technology, a generic tune MS method was used and the system was calibrated every five samples. The MS was operated in the positive full scan polarity mode (TOF start mass 250 Da; TOF stop mass 700 Da). The accurate mass of the monitoring ions ± 2 mDa was used for test item and internal standard peak integration. Full MS-TOF was applied with the m/Z ranges and mass resolutions of the Q-TOF set to ‘’high’’. Further analyzer settings were as follows: curtain gas 35, ion source gas1 50, ion source gas2 50, temperature 450°C, accumulation time 0.25 s, declustering potential 80.

**Table S6**. HPLC gradients (LC-MS/MS analysis)

| **7.44 compound** | | **Solubility / PPB / Plasma Stability** | | | | | |
|---|---|---|---|---|---|---|---|
| [min] | | 0.00 | 0.10 | 0.40 | 1.70 | 1.80 | 2.50 |
| Mobile phase | A (%)[a] | 0 | 0 | 97 | 97 | 0 | 0 |
| | B (%)[b] | 100 | 100 | 3 | 3 | 100 | 100 |
| **Propantheline** | | **Plasma Stability** | | | | | |
| [min] | | 0.00 | 0.10 | 0.40 | 1.70 | 1.80 | 2.50 |
| Mobile phase | A (%)[a] | 5 | 5 | 97 | 97 | 5 | 5 |
| | B (%)[a] | 95 | 95 | 3 | 3 | 95 | 95 |
| **7.44 compound / Omeprazole** | | **Chemical Stability & Hepatocyte Stability** | | | | | |
| [min] | | 0.00 | 0.10 | 0.40 | 2.70 | 2.80 | 5.50 |
| Mobile phase | A (%)[a] | 5 | 5 | 97 | 97 | 5 | 5 |
| | B (%)[a] | 95 | 95 | 3 | 3 | 95 | 95 |

[a] A: ACN / 0.1% (v:v) FA.

[b] B: $H_2O$ / 0.1% (v:v)FA.

**Table S7** gives an overview of the MS and chromatography parameters used for the analytes and the internal standard (ISTD).

**Table S7**. MS and chromatographic parameters (LC-MS/MS)

| **Triple quadrupole, ESI positive** | | | | | | |
|---|---|---|---|---|---|---|
| Compound | Molecular weight | $[M+H]^+$ (m/z) | Monitoring ion (m/z) | Scan time (s) | Collision energy (V) | RT (min) |
| **7.44** compound | 342.3 | 343 | 149 | 0.010 | 15 | 1.13 |
| Griseofulvin (ISTD) | 352.80 | 353 | 215 | 0.010 | 25 | 1.26 |
| Warfarin | 308.33 | 309 | 251 | 0.100 | 15 | 1.31 |

| | | | | | | |
|---|---|---|---|---|---|---|
| Propantheline | 448.39 | 368 | 181 | 0.015 | 35 | 0.95 |
| Diazepam (ISTD) | 284.7 | 285 | 193 | 0.015 | 20 | 1.19 |
| **LC-HRMS (QTOF)** | | | | | | |
| Compound | Molecular weight | $[M+H]^+$ (m/z) | Scan time (s) | | Collision energy (V) | RT (min) |
| **7.44** compound | 342.3 | 343.085 | 0.557 | | 10 | 2.10 |
| Omeprazole | 345.4 | 346.122 | 0.557 | | 10 | 1.85 |
| Diclofenac (ISTD) | 296.1 | 296.0239 | | | | 2. 09 |

**Fluorescence plate reader - Wallac Victor (Perkin Elmer)**

The fluorescent metabolites were detected using a Wallac Victor[3] fluorescence plate reader. The wavelengths for excitation and emission of the individual fluorescent metabolites depending on the substrates are given in **Table S8**. The final concentrations of the reagents required for the individual CYP isozymes are given. Shortcuts given in the tables are shown below.

**Table S8**. Excitation and emission wavelengths in P450 screening[a]

| Isoenzyme | CYP1A2 | CYP2B6 | CYP2C8 | CYP2C9 |
|---|---|---|---|---|
| Metabolite | CHC | HFC | F | HFC |
| Excitation | 405 nm | 405 nm | 485 | 405 nm |
| Emission | 460 nm | 535 nm | 545 | 535 nm |
| Isoenzyme | CYP2C19 | CYP2D6 | CYP3A4 | |
| Metabolite | CHC | AHMC | HFC | |
| Excitation | 405 nm | 380 nm | 405 nm | |
| Emission | 460 nm | 460 nm | 535 nm | |

[a]AHMC= 3-[2-(*N*,*N*-diethylamino)ethyl]-7-hydroxy-4-methylcoumarin, CHC = 3-cyano-7-hydroxycoumarin, F = fluorescein, HFC = 7-hydroxytrifluoromethyl coumarin

**Data analysis**

**Solubility in aqueous buffer**

The aqueous solubility (µM) of the compound was calculated using the following equation:

$$\text{Compound solubility } [\mu M] = \text{concentration in buffer supernatant } [\mu M] \qquad \text{S14}$$

**PPB**

The test item concentration in the 60 minutes stability control sample was compared to the non-incubated negative control sample concentration (= 100%) to proove test item plasma stability.

The percentage of compound bound to plasma proteins (% PPB) was calculated using the following equations:

(B) PPB referring to plasma filtrate:

$$\text{PPB [\%]} = 100 - \frac{(\text{test item concentration}_{\text{plasma filtrate}} / \text{RV})}{\text{mean test item concentration}_{\text{incubated plasma}}} * 100 \qquad \text{S15}$$

Results are expressed as mean PPB value calculated from plasma filtrate. In addition, the concentration of test item was corrected for the specific test item recovery (RV):

$$\text{RV} = \frac{(\text{concentration}_{\text{plasma filtrate}} [\text{nM}] + \text{concentration}_{\text{plasma retentate}} [\text{nM}]}{\text{mean concentration}_{\text{incubated plasma}} [\text{nM}]} \qquad \text{S16}$$

**Plasma stability and Chemical stability**

The amount of test item in the plasma stability samples and buffer (chemical stability) were expressed as percentage of remaining compound compared to time point zero (=100%). The depletion of test item was presented.

Half-life ($t_{1/2}$) estimates for the test item were determined using the rate of parent disappearance and following equation:

$$t_{1/2} = \frac{\ln 2}{-k} \qquad \text{S17}$$

$t_{1/2}$ = half life [min]

k = slope from the linear regression of log [test compound] versus time plot [1/min]

**Hepatocyte stability**

The amount of compound in the samples was expressed as percentage of remaining compound compared to time point zero (=100%). These percentages were plotted against the corresponding time points. *In vitro* intrinsic clearance ($CL_{int}$) and half-life ($t_{1/2}$) estimates were determined using the rate of precursor disappearance and following equation, based on the well-stirred liver model:

$$t_{1/2} = \frac{\ln 2}{-k} \quad \text{S18}$$

$t_{1/2}$ = half life [min]

k = slope from the linear regression of log [test compound] versus time plot [1/min]

$$CL_{int} = (-k) * V * f_u \quad \text{S19}$$

$CL_{int}$ = *in vitro* intrinsic clearance [µl/min/$10^6$ cells]

k = slope from the linear regression of log [test compound] versus time plot [1/min]

V = ratio of incubation volume and cell number

$f_u$ = unbound fraction in the blood

As $f_u$ is not known for the tested compound, the calculation was performed with $f_u = 1$.

$CL_{int}$ was used to calculate *in vivo* intrinsic clearance ($CL_{int}$ *in vivo*) based on S21. Scaling parameters are given in **Table S9.**

$$CL_{int}\ \textit{in vivo} = CL_{int} * w_{liver} * cd \quad \text{S20}$$

$CL_{int}$ *in vivo* = in vivo intrinsic clearance [ml/min/kg]

$CL_{int}$ = *in vitro* intrinsic clearance [ml/min/$10^6$ cells]

$w_{liver}$ = liver weight [g/kg]

cd = liver cell density [$10^6$ hepatocytes / g liver]

Hepatic clearance ($CL_{hep}$) was calculated as follows:

$$CL_{hep} = \frac{CL_{int}\ \textit{in vivo} * Q}{CL_{int}\ \textit{in vivo} + Q} \quad \text{S21}$$

$CL_{hep}$= hepatic clearance [ml/min/kg]
$CL_{int}$ *in vivo* = *in vivo* intrinsic clearance [ml/min/kg]
Q = blood flow [ml/min/kg]

**Table S9**. Scaling parameters for inter-species comparison

| **Species** | **Liver weight/ body weight [g/kg]** | **Liver blood flow [ml/min/kg]** | **Liver cell density [$10^6$ hepatocytes / g liver]** |
|---|---|---|---|
| Mouse (CD-1) | 87.5 | 90 | 135 |

**CYP inhibition using HTS™ kits**

Mean blank values (NC) were subtracted from the sample values to obtain the net fluorescence signals. For each inhibitor concentration, the percent inhibition was calculated relative to the wells without inhibitor (PC, no inhibition).

The resulting fluorescence signals of those compounds, for which auto-fluorescence has been detected, were corrected as follows. The mean values of the resulting auto-fluorescent signals of each test concentration (mean of three wells) were subtracted from the corresponding assay wells. The resulting corrected fluorescent signal was corrected by the blank value (NC).

### Permeability estimation from molecular dynamics simulations

#### System setup

To compute the permeability of **7.44**, a total of nine molecules with previously experimentally determined PAMPA measurements were selected as control compounds: progesterone, chlorpromazine, promazine, atropine, diazepam, theophylline, pralidoxime (2-PAM), asoxime (HI-6), and methoxime (MMB4). This set of molecules has been previously used to estimate permeabilities from MD simulations and free energy computations [13]. In all cases, atom types and their corresponding parameters were obtained from the AMBER GAFF2 force field [14], using antechamber. The restrained electrostatic potential (RESP) method was used to assign the charges from HF/6-31G calculations performed in Gaussian 09 [15]. Each simulation system was packed using PACKMOL-Memgen [16] to obtain a bilayer of 75x75 Å$^2$, resulting in ~165 1,2-dioleoyl-*sn*-glycero-3-phosphocholine (DOPC) molecules, 6585 TIP3P water molecules [17], and one ligand, in total ~43000 atoms. $K^+$ ions were added to neutralize the system when negatively charged compounds were simulated.

#### Molecular dynamics simulations

All MD simulations were performed using AMBER20 [14]. The minimization of the systems was performed using the MPI implementation of PMEMD [18]. 5000 cycles of steepest descent were followed in the first step of minimization by conjugate gradient minimization for a total of 10000 cycles. Only water molecules (and ions, if included) were minimized initially, using harmonic restraints of 25 kcal mol$^{-1}$ A$^{-2}$ on the rest of the system. In the second minimization step, the harmonic restraints were decreased to 5 kcal mol$^{-1}$ A$^{-2}$, while in the third and fourth steps of minimization, the restraints were applied only on the ligand. The fifth minimization step was performed without restraints. The minimized systems were then heated from 0 to 100 K for 5 ps in the NVT ensemble. The temperature was controlled using Langevin dynamics with a coupling constant of 1 ps$^{-1}$, SHAKE, and a time step of 2 fs in all cases [19]. Further heating to 300 K was performed under NPT conditions, using the Berendsen barostat with semiisotropic pressure scaling along the membrane plane for 115 ps. The simulations were further relaxed under the same conditions until 5 ns were obtained.

To calculate the permeabilities of the selected compounds, simulations were setup as previously demonstrated with the AMBER suite [14, 20]. In brief, simulations were carried out for each molecule, pulling for 32 ns at 300 K with a force constant of 1.1 kcal mol$^{-1}$ A$^{-2}$ and a pulling speed of 1 Å ns$^{-1}$ along the membrane normal (z-axis). A total of 33 umbrella windows were extracted, covering 0 to 32 Å with respect to the membrane center along the z-axis. In umbrella sampling simulations, each extracted structure was simulated for 100 ns, maintaining as a reaction coordinate the distance to the membrane center along the z-axis in the umbrella window with a force constant of 2.5 kcal $mol^{-1}$ $Å^{-2}$. The initial 50 ns of each simulation were considered as equilibration time, with the last 50 ns of each simulation being used for further analysis.

**Permeation potential of mean force and permeability calculations**

From the distance distributions obtained from the umbrella sampling simulations, PMF profiles were calculated using the weighted histogram analysis method (WHAM) [21], assigning the value of the probed molecule in bulk water to zero (**Figure 8A**). To calculate the membrane permeability (*P*, eq. S24), the diffusion along the membrane normal (*z*-axis) ($D(z)$, eq. S22), was calculated for each window as described by Hummer [22], implemented by Lee, Comer [23], and adapted by Dickson [20]:

$$D\big(z = (z)\big) = \frac{var(z)}{\tau_z} \quad \text{(S22)}$$

where $\tau_z$ corresponds to the characteristic time of the *z*-position autocorrelation in the given window. The diffusion $D(z)$ together with the free energy profile ($\Delta G(z)$) were used to obtain the resistivity ($R$) along the *z*-axis (eq. S23) and integrated to obtain the effective permeability $P_{eff}$ [23] (eq. S24):

$$R(z) = \frac{e^{\beta(\Delta G(z))}}{D(z)} \quad \text{(S23)}$$

$$P_{eff} = \frac{1}{R} = \frac{1}{\int_0^z R(z)\, dz} \quad \text{(S24)}$$

where β is the inverse of the Boltzmann constant times the absolute temperature, and the range 0 to $z$ covers the width of the whole membrane [13, 24]. Errors of the calculated permeabilities were estimated by performing the calculations in ten slices of 5 ns from the total 50 ns used for the analysis.

From the computed permeability data of our simulations and the experimentally determined PAMPA data of the reference molecules by Bennion *et al.* [13] ($\log P_{eff/PMF}\, P_0^{-1}$ and $\log P_{eff/PAMPA}\, P_0^{-1}$, respectively, **Table S10**), a linear regression was computed (Figure 8B). The calculated permeability rates ($\log P_{eff/PMF}\, P_0^{-1}$) of **7.44** in a protonated and deprotonated state were used to predict the experimental values ($\log P_{eff/PAMPA}\, P_0^{-1}$) from the linear regression (**Table S11**).

**Table S10**. Permeability data of reference compounds

| **Compound** | **log $P_{eff/PMF}$ $P_0^{-1}$ [a]** | **log $P_{eff/PAMPA}$ $P_0^{-1}$ [b]** |
|---|---|---|
| progesterone | 0.70±0.28 | -4.94 |
| theophylline | 0.12±0.50 | -5.91 |
| pralidoxime (P2-PAM) | -9.71± 0.74 | -7.52 |
| atropine | -0.75±0.94 | -5.26 |
| chlorpromazine | 1.58±0.18 | -5.26 |
| diazepam | 1.10±0.16 | -5.40 |
| asoxime (HI-6) | -14.43±0.91 | -11.16 |
| methoxime (MMB4) | -10.67±0.87 | -9.25 |
| promazine | 1.46±0.17 | -4.88 |

[a] $P_{eff/PMF}$, effective permeability in cm $sec^{-1}$ calculated from free energy calculations (eq. S24). $P_0$, unit factor corresponding to 1 cm $sec^{-1}$. Errors correspond to the standard deviation obtained from calculating the permeability when dividing 50 ns into ten independent 5 ns simulation slices.

[b] $P_{eff/PAMPA}$, effective permeability in cm $sec^{-1}$ obtained from PAMPA assays Bennion, Be [13]. $P_0$, unit factor corresponding to 1 cm $sec^{-1}$.

**Table S11**. Permeability data of **7.44.**

| **Compound** | **log $P_{eff/PMF}$ $P_0^{-1}$ [a]** | **log $P_{eff/PAMPA}$ $P_0^{-1}$ [b]** | **$P_{eff}$ [cm $sec^{-1}$]** |
|---|---|---|---|
| P7.44[c] | 0.89±0.34 | -5.07±0.10 | 8.51 x $10^{-6}$ |
| D7.44[d] | -1.90±0.87 | -5.82±0.24 | 1.51 x $10^{-6}$ |

[a] $P_{eff/PMF}$, effective permeability in cm $sec^{-1}$ calculated from free energy calculations (eq. S24). $P_0$, normalization factor corresponding to 1 cm $sec^{-1}$. Errors correspond to the standard deviation obtained from calculating the permeability from dividing 50 ns into ten independent 5 ns simulation slices.

[b] $P_{eff/PAMPA}$, effective permeability in cm $sec^{-1}$ calculated from the linear regression obtained from compounds in Table S10. $P_0$, normalization factor corresponding to 1 cm $sec^{-1}$.

[c] P7.44: **7.44** in the protonated (neutral) state;

[d] D7.44: **7.44** in the deprotonated (negatively charged) state.

### Supplemental Figures

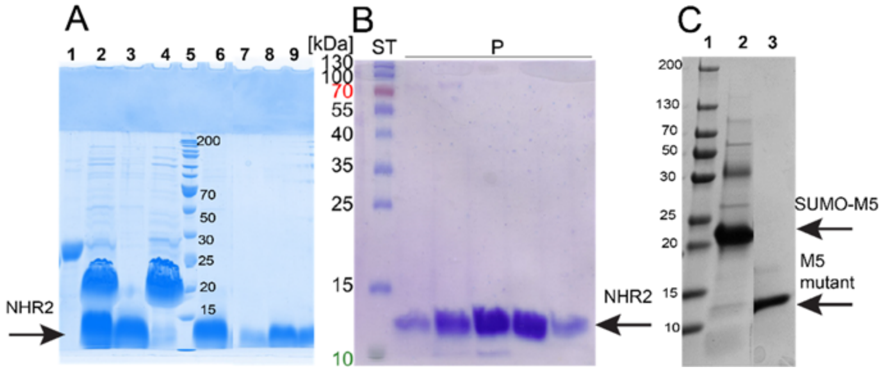

**Figure S1. SDS-PAGE of the NHR2 and M5 mutant purification.** The fractions from the elution step of the IMAC were analyzed using a 15% polyacrylamide gel. **A)** The purification of NHR2 from SUMO fusion protein. Lane numbers from left: 1- SUMO-NHR2 fusion protein from IMAC, 2- SUMO cleavage reaction, 3- NHR2 from a flowthrough from reverse IMAC, 4- SUMO elution from reverse IMAC, 5- PageRuler™ Marker, 6 to 9- SEC elution (8.5 kDa). **B)** Purification of 6His-NHR2. The protein bands correspond to the molecular weight of the NHR2 monomer (11.3 kDa). ST: 5 µl protein standard (PageRuler™ Prestained Protein Ladder; Thermo Scientific), P: fractions from the purification. **C)** The purification of M5 mutant from SUMO fusion protein. Lane numbers from left: 1-PageRuler™ Marker, 2- SUMO-M5 fusion protein from IMAC, 3- SEC elution (8.1 kDa).

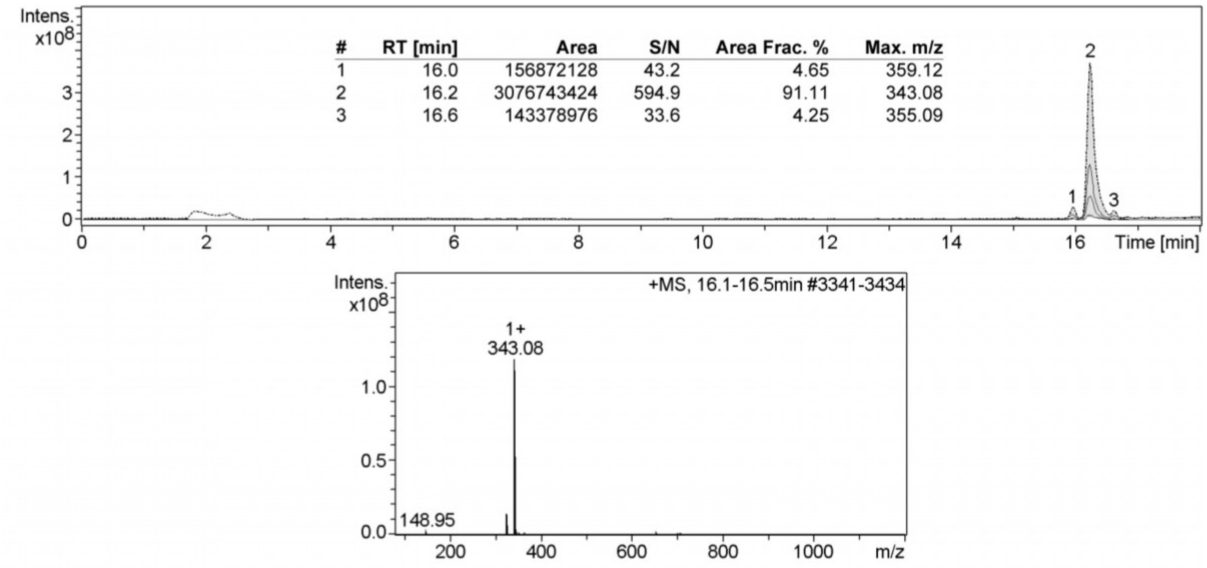

**Figure S2. LC-MS analysis of 7.44**. Ethanol-dissolved **7.44** (1 mg/mL) was used for LC-MS (amaZon speed ETD - Ion Trap, Bruker) analyses. The ionizations were generated by ESI. The upper panel is the chromatographic separation of **7.44** performed on a C18 reversed-phase analytical column. The most abundant compound **7.44** (peak 2) and lower amounts of other compounds (peak 1 and 3) were detected, and the respective masses are given as an insert. The lower panel is the MS data of **7.44** eluted from peak 2. The analyses of samples were carried out after three transitions for the sample. The first transition was for quantification and the second and third for verification.

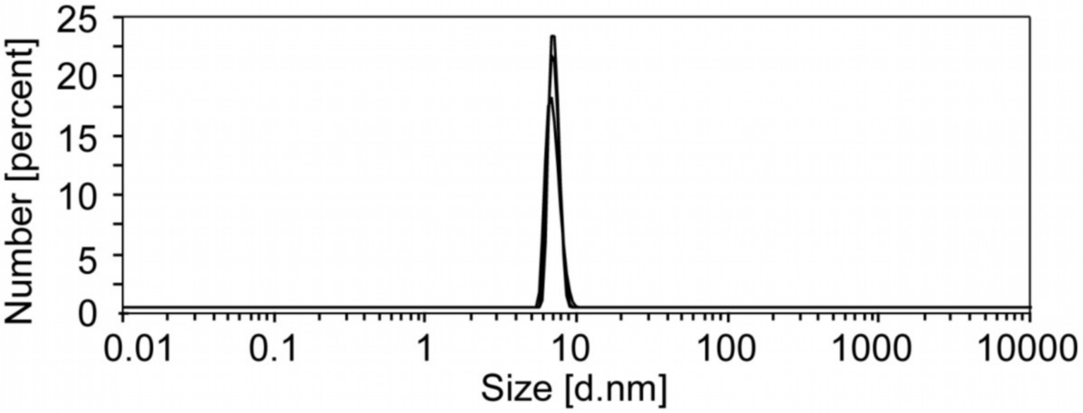

**Figure S3. Graph of the size distribution by number of NHR2 obtained from dynamic light scattering (DLS) measurements**. DLS experiments were performed at 298 K with a Zetasizer Nano S (Malvern Panalytical) light scattering instrument. The measurements were carried out with 53 µM and 25 µM of NHR2 protein in 20 mM sodium phosphate, 50 mM sodium chloride, 0.5 mM tris(2-carboxyethyl)phosphine, 10% (v/v) DMSO, pH 6.5. The plot of the size distribution versus the number shows a homogeneous distribution of the NHR2 sample with the average hydrodynamic radius of 6.939 nm.

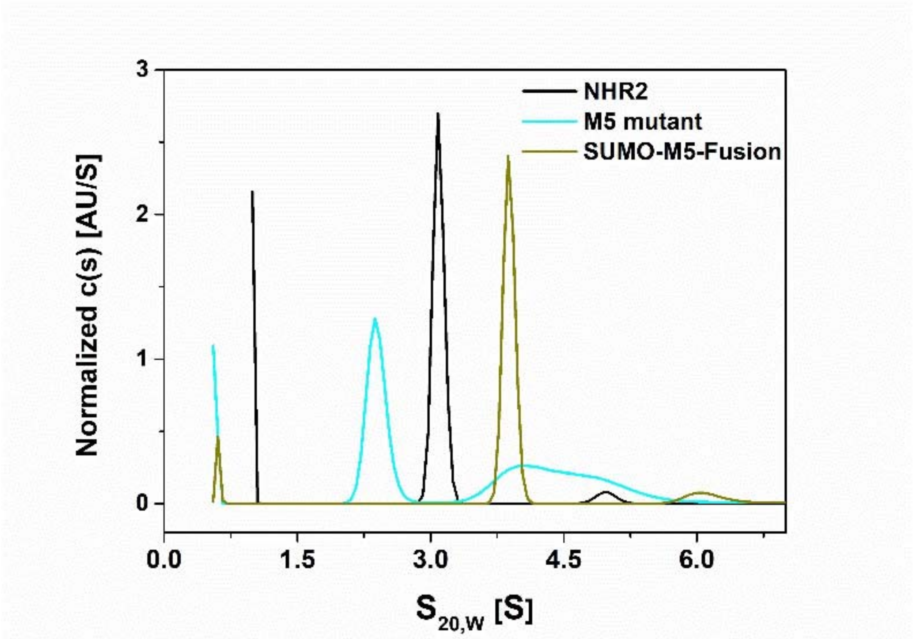

**Figure S4. Analysis of NHR2 oligomerization by analytical ultracentrifugation**. Sedimentation velocity concentration profiles of 13 µM of NHR2, 88 µM of M5 mutant, and 88 µM fusion protein of SUMO M5 mutant were obtained with a Beckman Coulter XL-I instrument in aluminum-filled double sector cells at 20°C and 50,000 RPM in 20 mM sodium phosphate, 50 mM sodium chloride, 0.5 mM tris(2-carboxyethyl)phosphine, pH 6.5. Samples were measured at 280 nm. The sedimentation profiles were fit to finite element solutions of the Lamm equation using a model of a continuous distribution of discrete, noninteracting species with the program SEDFIT [25]. The measured sedimentation coefficient (S) values were 3.07, 2.28, 3.89 for NHR2, M5 mutant, and SUMO-M5 fusion protein, respectively. The measured molecular weights were 37.3 kDa, 28.0 kDa, and 43.4 kDa, and the theoretically calculated molecular weights are 34.2 kDa (tetramer), 17.1 kDa (dimer), and 43.9 kDa (dimer) for NHR2, M5 mutant, and SUMO-M5 fusion protein, respectively. The measured molecular weight of the M5 mutant is higher than the calculated molecular weight. Hence, to clarify the size of the dimer, the SUMO-M5 fusion protein was measured, and the expected molecular weight of the dimer was obtained. The molecular weight and sedimentation coefficient of the predominant species were determined using a continuous distribution model (c(s)) [26]. A weight percent contribution of 94% of the majority species (tetramer) was obtained for 35 µM of NHR2 by integration of a continuous sedimentation coefficient distribution function. Then, the same 35 µM concentration of NHR2 was used for the SEC study (see main text about SEC and Figure 2A).

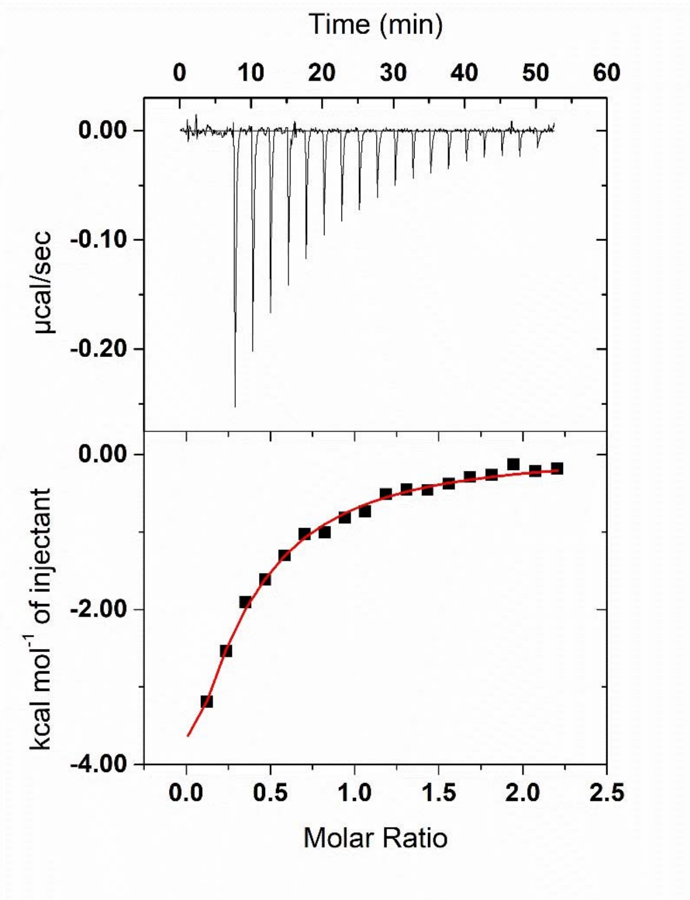

**Figure S5. NHR2 interaction with E proteins (HEB (176-200)) by isothermal titration calorimetry (ITC)**. Titration of 600 µM HEB (176-200) peptide into a 53 µM NHR2 solution and resulting heat change are shown in the upper panel. The integrated data were fitted to a one-site binding model (lower panel) to yield $K_D = 30 \pm 4$ µM and $\Delta H = -15 \pm 6$ kcal $mol^{-1}$. ITC measurements were performed at 25°C using a MicroCal ITC200 (Malvern Instruments Ltd., United Kingdom) calorimeter. Buffer conditions were 50 mM sodium phosphate, 50 mM sodium chloride pH 8. Data were fitted using Origin Software package (OriginLab, Northampton, MA) provided by Malvern company.

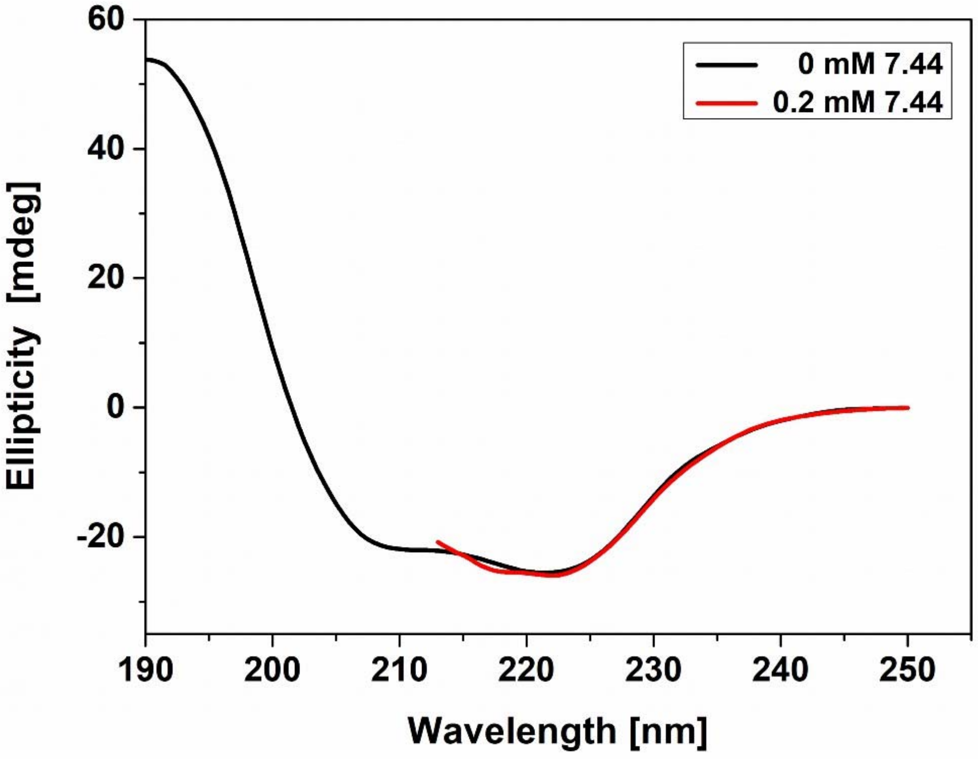

**Figure S6. CD spectrum of NHR2 apo and in complex with 7.44**. CD spectra were measured at 25 °C for the sample of 12.5 µM NHR2 for the apo state (black) and in complex with 200 µM of **7.44** (red). The samples were diluted in 10 mM sodium phosphate, 10 mM sodium chloride, 0.5 mM tris(2-carboxyethyl)phosphine, <1% (v/v) DMSO pH 6.5. Due to the scattering (high voltage in CD) of DMSO in the complex sample, the spectrum was measured from 213 nm to 250 nm, whereas the apo protein was measured from 190-250 nm.

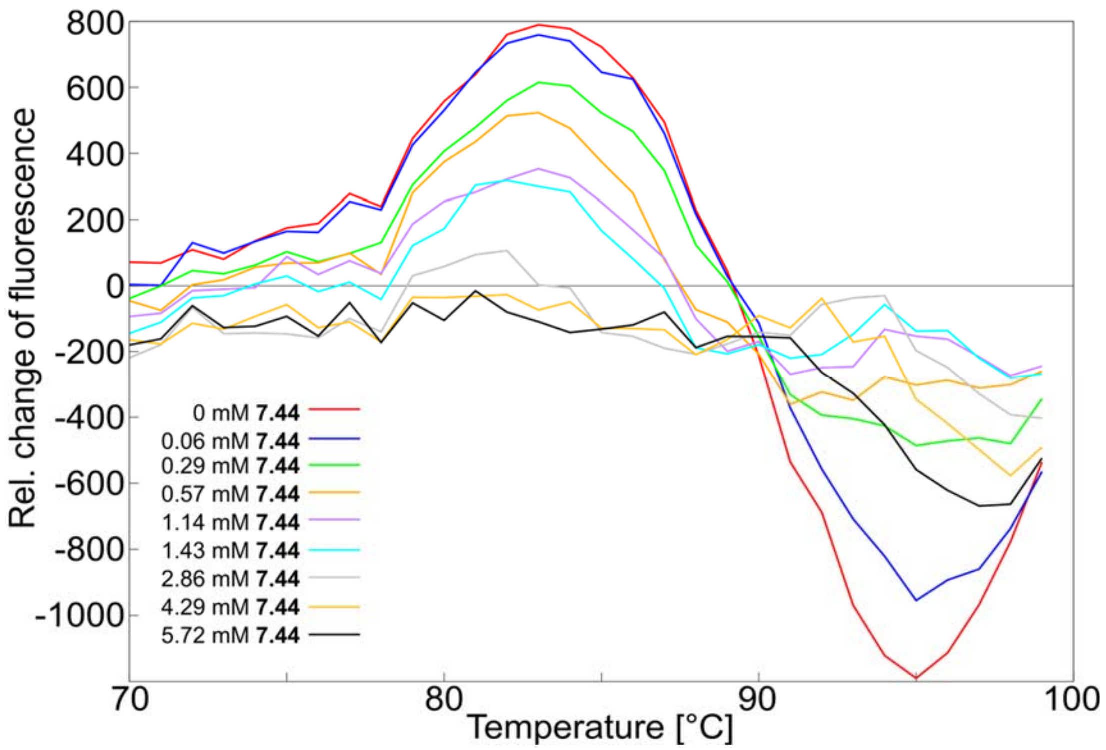

**Figure S7. Differential scanning fluorimetry**. dF/dT was calculated for nine samples containing a constant concentration of NHR2 and a varying concentration (provided in the legend) of **7.44**.

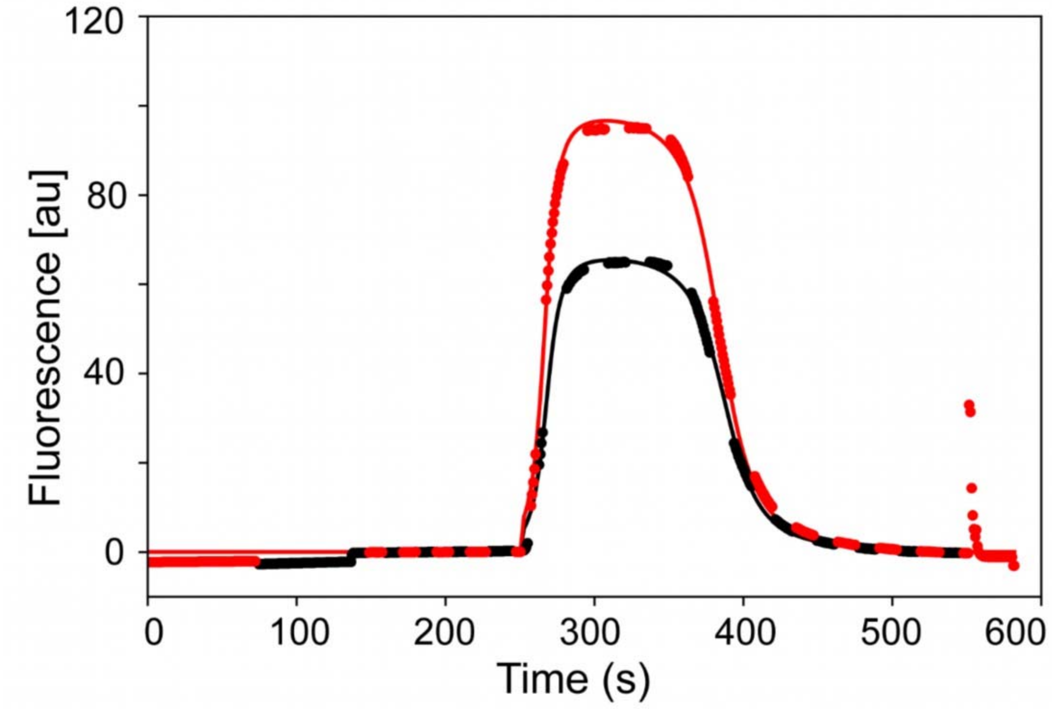

**Figure S8. Measurement of the hydrodynamic radius of fluorescently labeled NHR2 by using microfluidic diffusional sizing method**. Hydrodynamic radius was measured for 100 nM concentration of Alexa 488 dye-labeled NHR2 sample in 50 mM sodium phosphate, 50 mM sodium chloride pH 8, with microfluidic diffusional sizing using the Fluidity One-W instrument (Fluidic Analytics Ltd) at 25 °C. Fluorescence emission intensity at ~520 nm of labeled NHR2 after injection into a microfluidic laminar flow chamber and separation into two detection channels corresponding to diffused (black) and undiffused fluorophores (red). The measured hydrodynamic radius was 2.22 nm, which corresponds to the molecular weight of 16.72 kDa (NHR2-dimer) calculated by the software supplied with the machine [27]. Experiments were performed as triplicates.

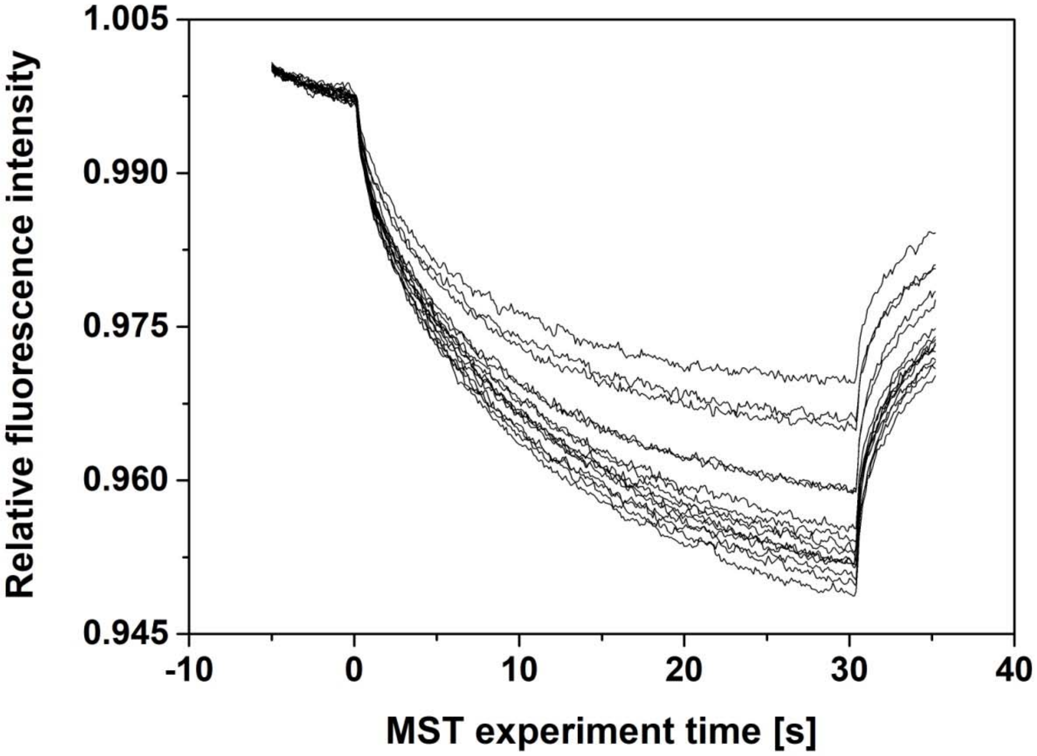

**Figure S9**. **MST traces of 7.44 binding to NHR2.** 100 nM of labeled NHR2 was mixed with **7.44** to concentrations ranging from 122 nM to 4 mM to the final volume of 50 µl (serial 1:1 dilution) in 20 mM sodium phosphate, 50 mM sodium chloride, pH 8.0, 10% (v/v) DMSO. The samples were incubated overnight in the dark prior to the experiment. Thermophoresis was measured using a Monolith NT.115 instrument, using an excitation power of 50% for 30 s and MST power of 40% at an ambient temperature of 24 °C. Microscale thermophoresis results were analyzed by MO affinity analysis software (NanoTemper Technologies GmbH).

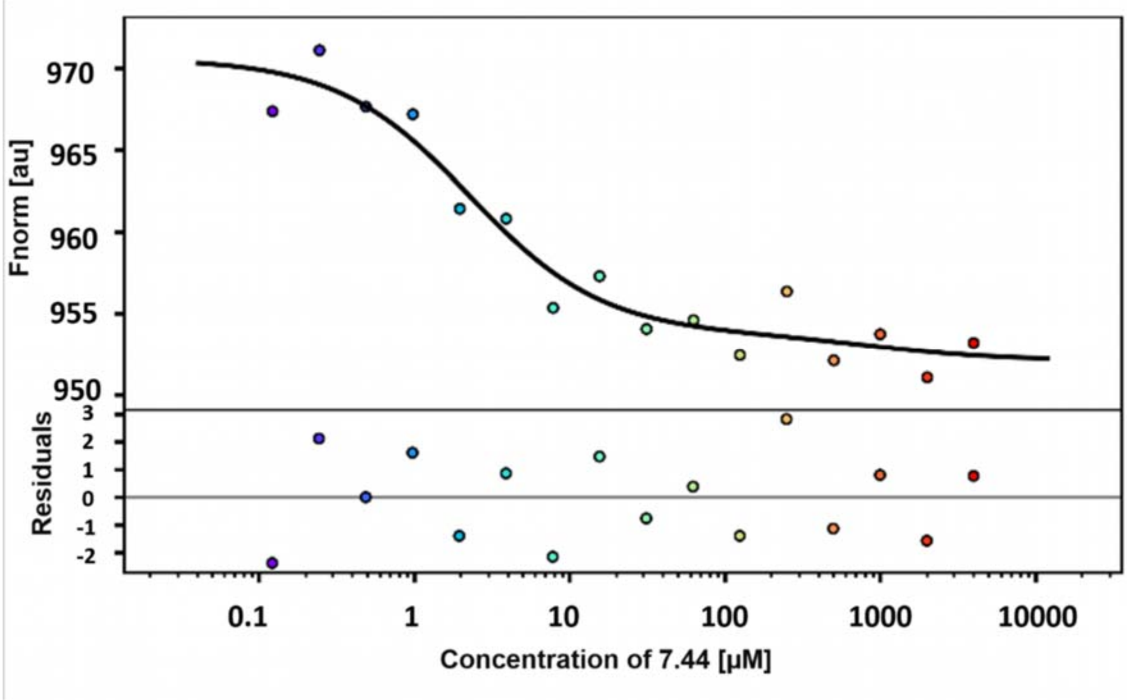

**Figure S10. MST studies of 7.44 binding to NHR2.** Titration of **7.44** to a constant concentration of dye-labeled NHR2 induces a change in thermophoresis. Data were fitted to a 1:2 binding model, assuming a 2-fold symmetry in the NHR2 dimer using the PALMIST software [28, 29]. The fitting results in $K_{lig}$ = 2 µM, 68.3% CI [0, 6] µM. The broad confidence interval is a result of too few data points to clearly resolve a second binding step due to the insolubility of **7.44** at higher concentrations.

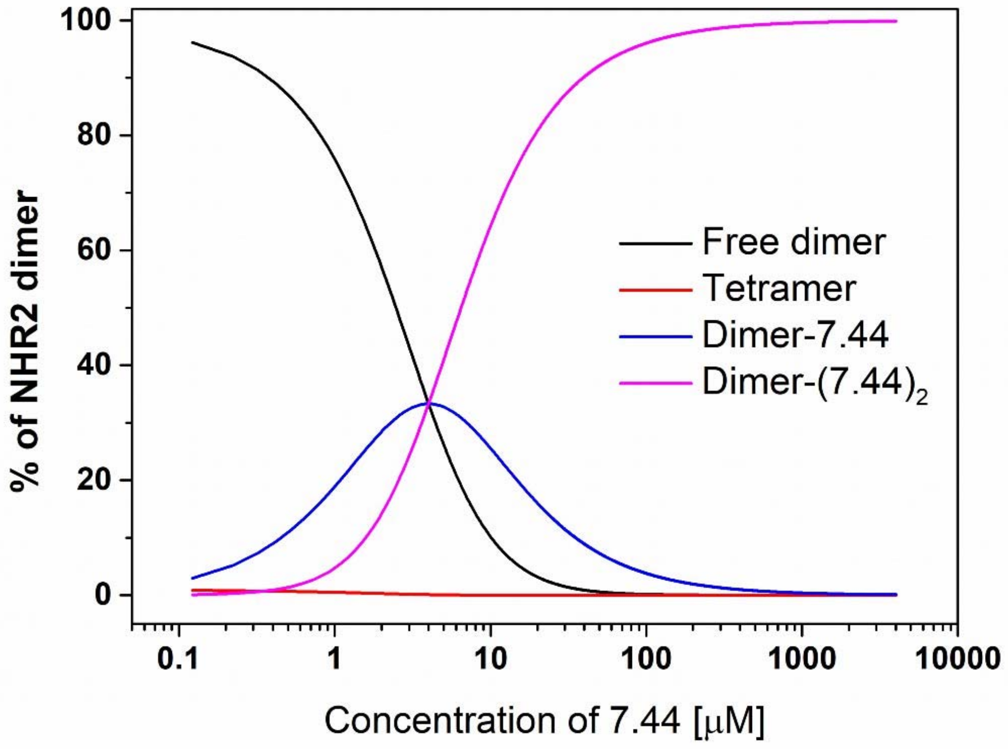

**Figure S11. Simulated species distributions of the NHR2 / 7.44 equilibrium (eq. 4 in the main text)**. The distributions for NHR2 tetramer and dimer as well as singly and doubly **7.44**-bound dimer were obtained from the determined $K_{tet}$ and $K_{lig}$ using HySS2009 [30]. In the absence of **7.44**, > 99% of NHR2 is in a dimer configuration under the experimental conditions of the MST experiments. Increasing the **7.44** concentration reduces the amount of free NHR2, including the NHR2 tetramer. For calculating $K_{lig}$ according to eqs. S11 and S12, it was assumed that the majority of the MST signal change is attributable to the dimer, yielding that, at $EC_{50}$ = 2.5 μM of **7.44** (eq. 3 in the main text), the amount of free NHR2 dimer in the absence of **7.44** is reduced by half. This is retrieved back from the simulation.

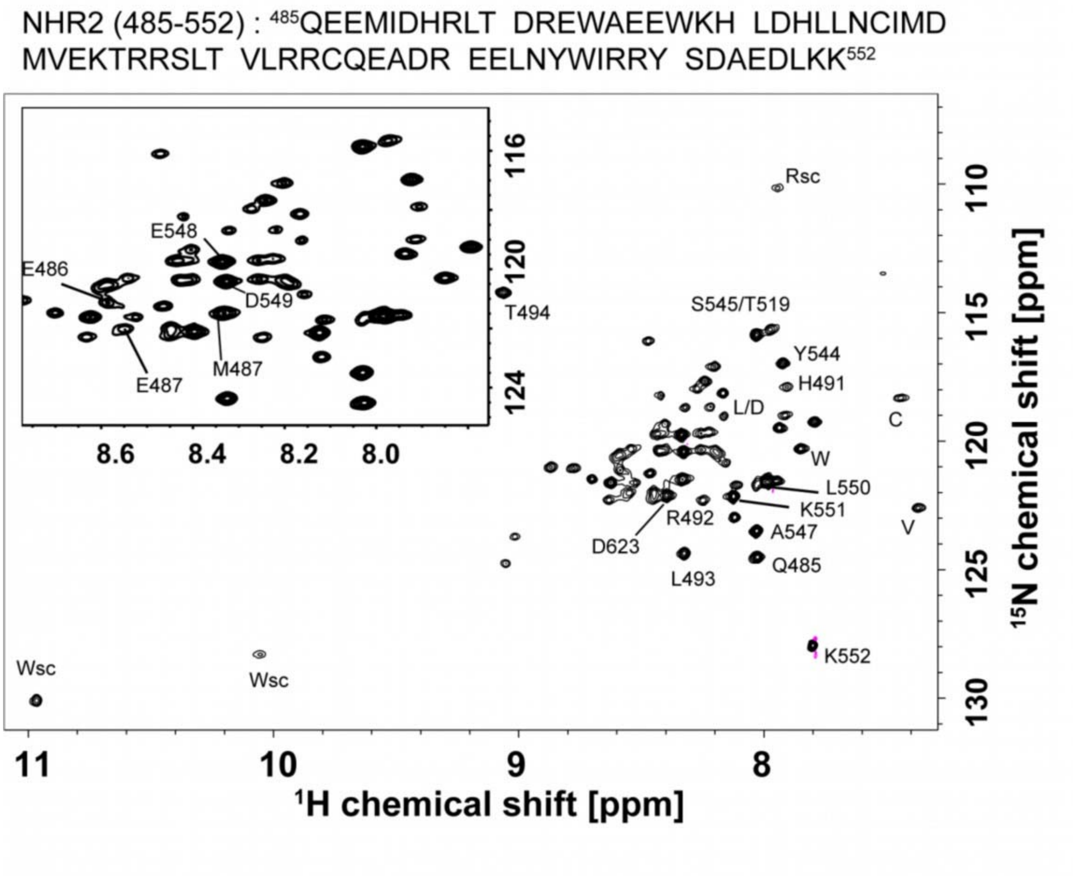

**Figure S12. 2D $^{1}$H-$^{15}$N TROSY-HSQC spectrum of 47 µ M of uniformly $^{2}$H, $^{15}$N, $^{13}$C-labeled NHR2**. The amino acid sequence of NHR2 is depicted at the top of the spectrum. The spectrum was recorded at 60 °C on a 750 MHz spectrometer for about 19 hours. The assignments were done by 3D NMR experiments with 325 µM of $^{2}$H, $^{13}$C, $^{15}$N-NHR2 in 20 mM sodium phosphate, 50 mM sodium chloride, 0.5 mM tris(2-carboxyethyl)phosphine, 10% (v/v) $D_2O$, pH 6.5 in a Shigemi tube and at 308 K. 32% of the amide signals were unambiguously assigned and labeled in the spectrum. The assignments were hindered by missing signals in the 3D spectra, which is due to the coiled-coil nature of the protein.

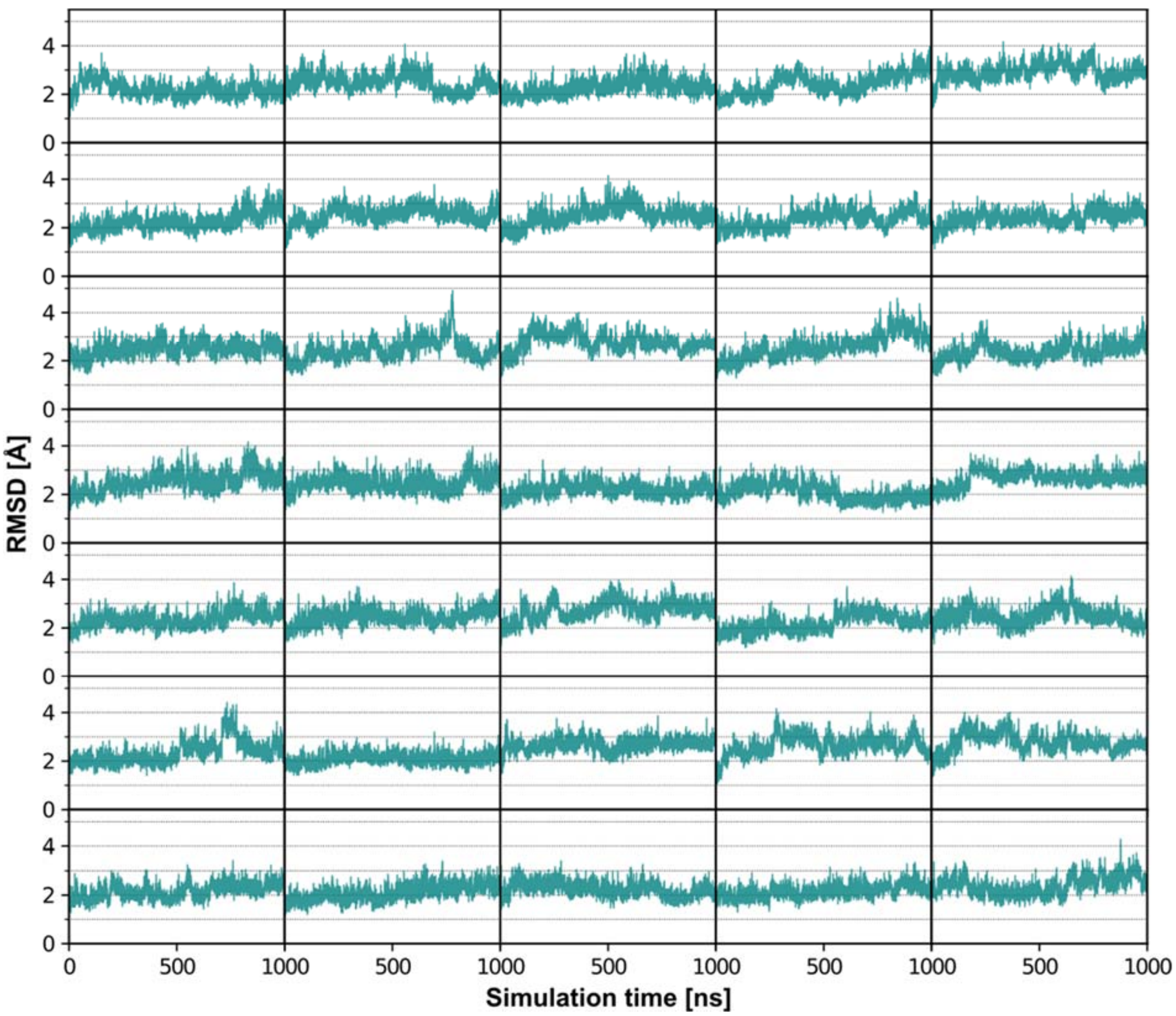

**Figure S13. Structural variability of the NHR2 tetramer in the molecular dynamics simulations.** The $C_\alpha$-RMSD of the NHR2 tetramer during the simulations with respect to the crystal structure is shown (PDB ID: 1wq6). Occasional higher RMSD values up to 5 Å correspond to motions of the dynamic termini, while the core structure of the tetramer remains stable.
